# RpS12-mediated induction of the Xrp1^short^ isoform links ribosomal protein mutations to cell competition

**DOI:** 10.1101/2025.06.15.659587

**Authors:** Myrto Potiri, Eleni Tsakiri, Kyriaki Kontogiannidi, Maria Loizou, Efthimios M.C. Skoulakis, Martina Samiotaki, Panagiota Kafasla, Marianthi Kiparaki

## Abstract

Cell competition, a universal yet enigmatic phenomenon, eliminates less-fit cells via interactions with their neighbors. It was originally described in *Drosophila* mosaics, where heterozygous ribosomal protein *(Rp*^+/−^) mutant cells are eliminated by wild-type neighbors. The transcription factor Xrp1 mediates most of the *Rp^+/−^* -associated phenotypes, including reduced competitiveness and translation. Although RpS12 is required for Xrp1 induction in *Rp*^+/−^ cells, the mechanism remained unresolved. We demonstrate that RpS12, via alternative splicing, induces the Xrp1 short (Xrp1^short^) isoform expression in *Rp^+/−^* cells, which is both necessary and sufficient for their elimination. Strikingly, RpS12 overexpression in wild-type cells is sufficient to induce Xrp1^short^ expression and confer a “loser” phenotype. While Xrp1^long^ isoform is not required in *Rp^+/−^* cells, expression of either Xrp1 isoform is sufficient to promote the loser status in wild-type cells. We further identify Syncrip, an RNA-binding protein reduced in *Rp*^+/−^ cells, as a critical Xrp1 suppressor; its depletion in wild-type cells activates Xrp1-dependent competition. Our findings establish RpS12’s specialized function in Xrp1^short^ promotion, not proteotoxic stress, as the primary driver in *Rp*^+/−^ cells, providing new perspectives that challenge and refine prevailing models. Our work contributes in long-standing questions about ribosomal protein-linked fitness surveillance and provides insights into ribosomopathy pathologies.

## INTRODUCTION

Life is inextricably linked to the function of ribosomal protein (*Rp*) genes, rendering most homozygous *Rp* mutations lethal. In humans, heterozygous mutations in *Rp* genes cause diseases such as ribosomopathies[1, 2] and cancer[3, 4]. In *Drosophila,* heterozygous dominant *Rp* mutants (*Rp*^+/−^) are viable and exhibit the “*Minute*” phenotype, characterized by slow growth, developmental delay and short bristles in the adult[5, 6].

However, in larval mosaic imaginal discs (epithelial structures that grow rapidly in an undifferentiated state in the larva and give rise to adult tissues upon metamorphosis), *Rp*^+/−^ cells are actively eliminated via apoptosis by wild-type neighboring cells[6–10]. *Rp*^+/−^ competitive apoptosis is predominantly localized at the boundaries next to wild-type cells (boundary cell death) and is the major hallmark of *Rp*^+/−^ driven cell competition[7]; consistent with the short-range nature of the phenomenon[11]. This was the first paradigm of cell competition, where, in mosaics, otherwise viable cells (referred as losers) are eliminated by neighboring cells (called winners) based on relative fitness differences, rather than intrinsic defects[6, 9, 10]. A plethora of cell competition examples have since been identified in both *Drosophila* and mammals, with different triggers and elimination mechanisms[12–15]. Cell competition is a highly dynamic process recognized in diverse biological contexts, including embryonic development, tissue homeostasis, ageing, and multiple diseases, such as cancer [10, 12, 14–20]. Nevertheless, the mechanisms driving cell competition remain incompletely understood.

Xrp1, a rapidly evolving AT-hook/bZip-domain transcription factor mediates most *Rp^+/−^*responses in *Drosophila*[21]. These Xrp1-dependent *Rp^+/−^* or “*Minute*” phenotypes include reduced cellular competitiveness in mosaics, slower cellular growth, altered gene expression, developmental delay, and reduced translational activity[9, 21–23]. Xrp1 function depends on its heterodimer partner Irbp18 (a ubiquitous bZip transcription factor) and both are upregulated in *Rp*^+/−^ cells via transcriptional autoregulation, which is essential for *Rp*^+/−^ responses[21, 22, 24–26]. Notably, heterozygous *Xrp1* null mutations (*Xrp1*^+/−^) are dominant haploinsufficient; heterozygous *Xrp1* mutations block Xrp1 activation in *Rp*^+/−^ cells and rescue *Minute* phenotypes[21, 25, 27].

The eukaryotic-specific ribosomal protein RpS12 (eS12), part of the ribosome’s beak region and encoded by a non-*Minute* gene [5], is required for Xrp1 activation in *Rp*^+/−^ cells [21, 27–29]. *Rp*^+/−^ cells homozygous for the *rpS12^G97D^* mutation, where glycine 97 is substituted by aspartic acid, fail to activate the Xrp1 pathway[21, 27, 28, 30]. Notably, this G97-dependent function of RpS12 is distinct from its essential role in cell growth and survival, as *rpS12^G97D^*homozygous flies are viable[27, 28]. While increasing RpS12 levels in *Rp*^+/−^ cells reduced their growth, this effect was not statistically significant in wild-type cells, strongly suggesting that RpS12 requires an additional signal from *Rp^+/−^* cells to activate Xrp1 and initiate competition[21, 27, 28].

In addition, Xrp1 has a crucial role as a stress sensor and effector, activated by a broad spectrum of stresses, including defects in ribosome biogenesis or function, endoplasmic reticulum (ER) stress, oxidative stress, proteasome impairment and spliceosome defects[9, 25, 31–37]. Beyond its role in cell competition, the RpS12-Xrp1 pathway coordinates inter-organ growth via Dilp8 regulation, ensuring tissue integrity during development[38]. Interestingly, this pathway also contributes to the elimination of aneuploid cells during *Drosophila* development[39]. Therefore, understanding Xrp1 activation by RpS12 is critical for deciphering diverse homeostatic pathways during development.

The mechanism of Xrp1 regulation by RpS12 has remained enigmatic, particularly whether RpS12 acts as the primary signal inducing Xrp1 in *Rp*^+/−^ cells, or requires additional *Rp*^+/−^specific signals. An alternative model proposed that proteotoxic stress, rather than RpS12, as the initial trigger of Xrp1 activation in *Rp*^+/−^ cells[31, 40, 41]. This stress was proposed to further induce Xrp1 activation in *Rp^+/−^* cells through a feed-forward-loop[31, 40, 41]. While *Rp*^+/−^ cells exhibit increased proteotoxic stress, reducing phosphorylated eIF2a (p-eIF2a) levels in *Rp*^+/−^ cells did not affect Xrp1 expression, whereas Xrp1 depletion abolished all proteotoxic stress markers (such as p62 and ubiquitin accumulation and elevated p-eIF2α)[9, 25, 32, 40, 41]. Therefore, our data supported that although proteotoxic stress can activate Xrp1 and competition in other contexts[25, 31, 32, 36], in *Rp*^+/−^ cells Xrp1 drives-rather than responds to-proteotoxic stress[9, 25]. Furthermore, despite the Xrp1’s requirement in various competitive contexts, it remained unknown whether Xrp1 is sufficient to initiate cell competition, or if it merely enhances or contributes to another instigator factor.

Here, we demonstrate that RpS12 overexpression in otherwise wild-type cells is sufficient to induce Xrp1 expression and trigger their competitive elimination in mosaic tissues, independent of ribosomal haploinsufficiency. Mechanistically, we show that RpS12 regulates the Xrp1 alternative splicing, leading to the production of an mRNA encoding the Xrp1 short protein isoform in both wild-type and *Rp*^+/−^ cells. Importantly, we prove for the first time that overexpression of either Xrp1 isoform is sufficient to induce loser status in wild-type cells. Accordingly, we show that the Xrp1^long^ isoform is not required for *Rp*^+/−^ competition, even if it is sufficient to induce competition in wild-type cells. Furthermore, we find that *Rp*^+/−^ cells exhibit reduced levels of the RNA-binding protein Syncrip, and that Syncrip depletion in otherwise wild-type cells is sufficient to induce their competitive elimination through Xrp1 activation. Together, these findings reveal a novel mechanism of Xrp1 activation via RpS12-dependent alternative splicing and offer mechanistic insights into how ribosomal protein mutations drive cell competition, with implications for understanding ribosomopathies.

## RESULTS

### RpS12 overexpression is sufficient to induce Xrp1-dependent cell competition

We unexpectedly found that RpS12 overexpression in wild-type disc clones was sufficient to induce Xrp1 expression (Figure 1A). These clones exhibited apoptotic cell death, predominantly at clone boundaries adjacent to wild-type cells (Figures 1B and 1C). Xrp1 depletion abolished this RpS12-dependent cell death, confirming Xrp1’s essential role in this process (Figure 1D).

**Figure 1.**
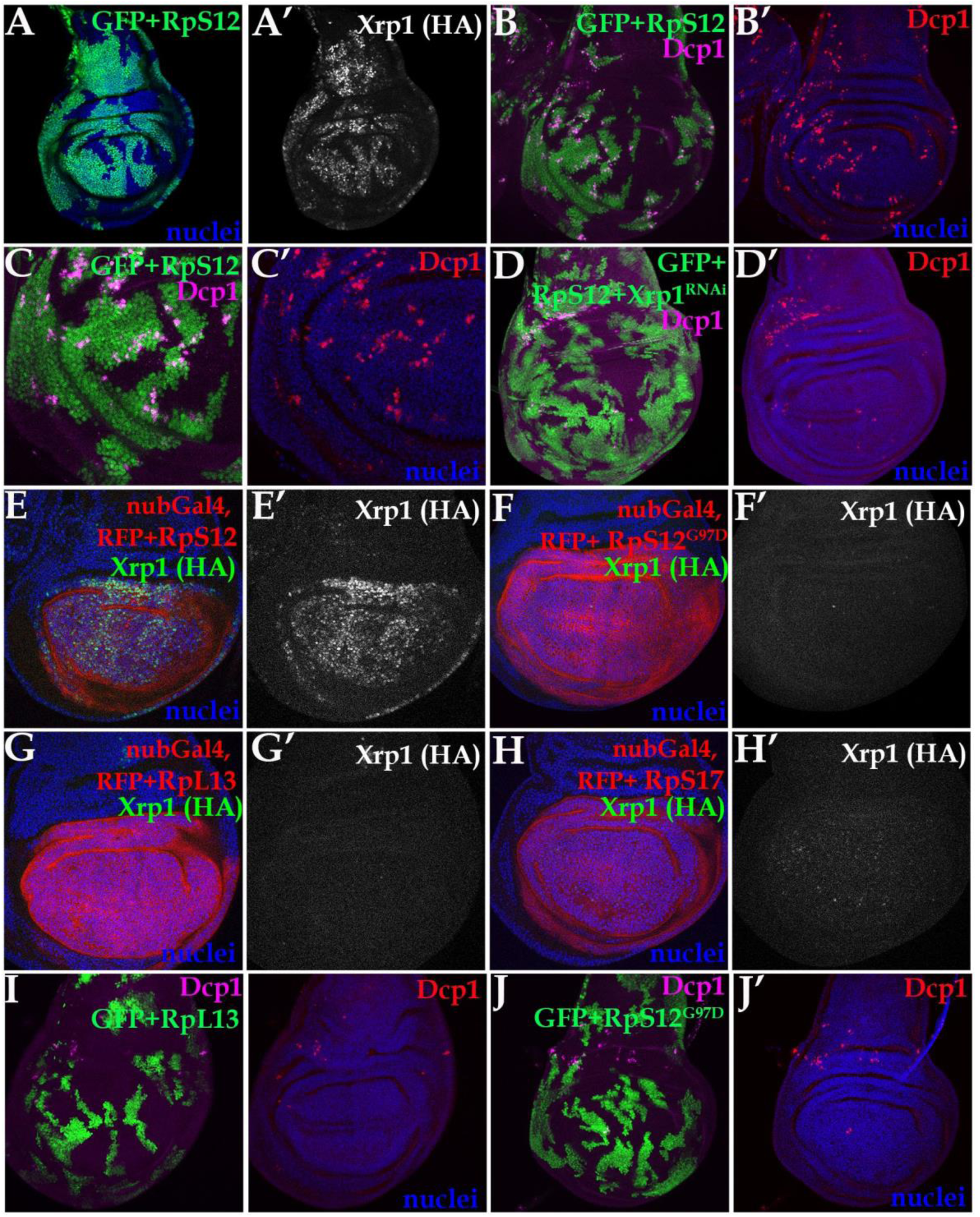
RpS12 overexpression is sufficient to induce Xrp1-dependent cell competition in wild-type cells. Confocal images of wing imaginal discs of the indicated genotypes. Panels (B)-(D), (I) and (J) show a projection of active Dcp1 staining of the central disc-proper region. Panels (A) and (E)-(H) show single confocal planes. (A) RpS12 overexpression in clones (green, GFP) increased the levels of the endogenously HA-tagged Xrp1 allele. (B, C) RpS12 overexpression in clones (green, GFP) in wild-type discs induced apoptotic cell death (Dcp1, magenta in B,C and red in B’, C’) at the boundaries with the wild-type cells (zoom panel in C). (D) RpS12-dependent boundary cell death (Dcp1, magenta in D and red in D’) in clones was completely abolished upon depletion of Xrp1 by RNAi (green, GFP). (E-H) Xrp1-HA expression from wing discs where nub-Gal4 drives the co-expression of UAS-RFP reporter (red) together with the indicated UAS transgenes. In (E), overexpression of RpS12 resulted in strong Xrp1-HA expression (E’, green in E). Overexpression of RpS12^G97D^ (F) or RpL13 (G) could not induce Xrp1-HA expression (F’-G’, green in F, G). In (H), overexpression of the RpS17 transgene induced negligible levels of Xrp1-HA expression (H’, green in H). (I-J) Clonal overexpression (green, GFP) in wild type discs of RpL13 (I) or RpS12^G97D^ (J) did not show increased cell death (magenta I, J and red in I’, J’). See also Figure S1

Prior studies in *Drosophila Rp^+/−^*mutants proposed that, akin to yeast, the accumulation of unassembled ribosomal proteins (RPs) overloads and suppresses the proteasome and autophagy systems, leading to protein aggregation and proteotoxic stress[31, 40–43]. To investigate whether RpS12 overexpression causes Xrp1-dependent cell competition through proteostasis collapse due to unassembled RPs, we examined the effects of overexpressing other ribosomal proteins. While RpS12 overexpression induced Xrp1 expression in wild-type discs, overexpression of other RPs, including the RpS12^G97D^ mutant, RpL14 or RpL13 did not induce Xrp1 expression, whereas RpS17 showed only minimal levels of Xrp1 (Figures 1E-1H, and S1A). Importantly, clonal overexpression of RpS12^G97D^, RpS17, RpL14 or RpL13 failed to trigger cell competition (Figures 1I, 1J, and S1B-S1C). Unexpectedly, we were unable to recover any clones upon overexpression of GFP-RpL10Ab protein, using a transgene employed for cell-specific profiling of translated mRNAs in *Drosophila* (Figure S1D)[44]. Intriguingly, overexpression of GFP-RpL10A induced higher Xrp1 levels compared to RpS12 overexpression, and reduced the compartment size, although the effects of untagged RpL10A remain to be determined (Figures S1E-S1F). Therefore, RpS12 driven cell competition presented striking specificity, dependent on its Gly97 dependent function and on Xrp1 induction.

### RpS12 regulates Xrp1 alternative splicing, producing a short Xrp1 isoform

The Xrp1 locus produces seven mRNA isoforms through differential promoter usage and alternative splicing[45]. Four transcripts (RA, RB, RF and RG) encode the 73kDa Xrp1 long protein isoform (Xrp1^long^), and the remaining three (RC, RD and RE) the 45kDa short isoform (Xrp1^short^) (Figure 2A)[45]. At their C termini, both protein isoforms contain the AT-hook and the b-ZIP DNA binding motifs, whereas their N-termini differ. Exon 4 contains the translation start site (ATG) for the Xrp1^long^ isoform, while the ATG for the Xrp1^short^ isoform is located on exon 5 (Figures 2A and S3A).

**Figure 2.**
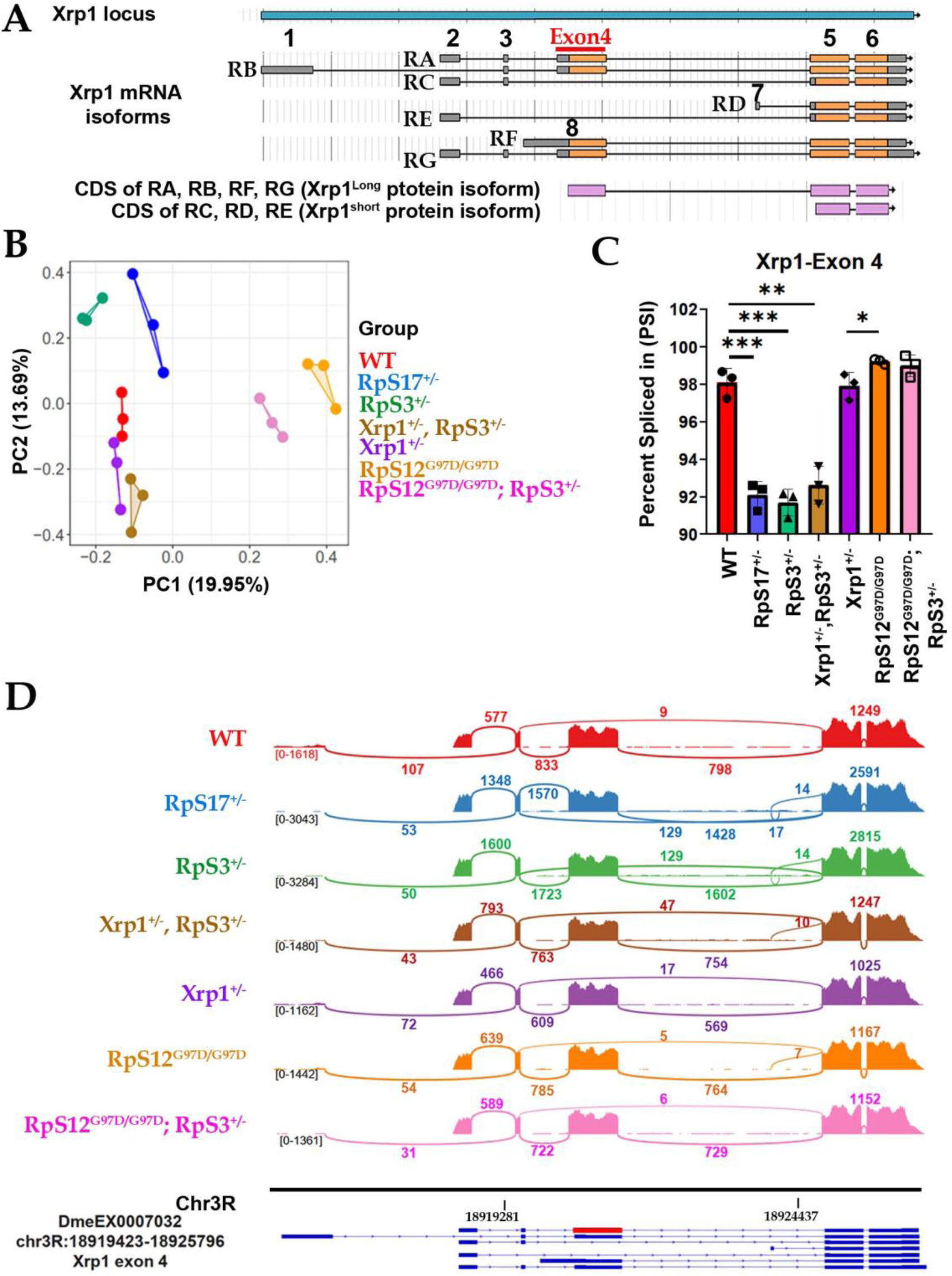
RpS12 promotes exon 4 skipping of Xrp1 mRNA, producing the Xrp1^short^ isoform. (A) 7 different isoforms are predicted to be transcribed from the Xrp1 locus, four of them (RA, RC, RE and RG) having the same transcription start site. RA, RB, RF and RG encode the long protein isoform of Xrp1, while the RC, RD and RE encode the short Xrp1 isoform. (B) Principal Component Analysis (PCA) illustrates the distribution of the PSI values for all exonic splicing events mapped across all the genotypes depicted. Each point represents an individual sample analyzed. (C) PSI values of the *Xrp1* exon 4 inclusion in the final mRNA, represented as mean ± SEM. *p-values* are calculated by Student’s t-test (** p< 0.01, *** p< 0.001). (different genotypes presented: wild-type in red, *RpS17*^+/−^ in blue, *RpS3*^+/−^ in green, *Xrp1^+/−^, RpS3^+/−^*in brown, *Xrp1^+/−^* in purple, *RpS12^G97D^*^/G97D^ in orange and *RpS12^G97D^/^G97D^,RpS3^+/−^* in pink) (D) Sashimi plots for visualization of the *Xrp1* alternative splicing events derived from aligned RNAseq data. Numbers of exon-exon junction reads are indicated above or below the arcs. At the bottom, a schematic shows a view of the exons and the introns of the different *Xrp1* mRNA isoforms. The chromosomal location is indicated below. The box in red indicates exon 4 of the *Xrp1*, with coordinates Chr3R: 18919423-18925796 and the VastDB identification number DmeEX0007032. (different genotypes presented: wild-type in red, *RpS17*^+/−^ in blue, *RpS3*^+/−^ in green, *Xrp1^+/−^, RpS3^+/−^* in brown, *Xrp1*^+/−^ in purple, *RpS12^G97D/G97D^* in orange and *RpS12^G97D^/^G97D^,RpS3^+/−^*in pink) See also Figure S2

Previous research identified Xrp1 mRNA as a highly occupancy RNA target, interacting with 17 of 20 RNA binding proteins (RBP) studied, including multiple splicing factors[46]. Additionally, Xrp1 is efficiently co-transcriptionally spliced in *Drosophila* S2 cells[47]. To investigate splicing-mediated regulation of Xrp1, we performed vast-tools analysis on our previously published RNA-seq datasets[21, 27] to profile changes in alternative splicing (AS) pattern across seven genotypes: a) wild-type; b) *RpS17*^+/−^; c) *RpS3*^+/−^; d) *Xrp1*^+/−^, *RpS3*^+/−^; e) *Xrp1*^+/−^ ; f) *RpS12^G97D^*^/G97D^; and g) *RpS12^G97D^*^/G97D^ *,RpS3*^+/−^ (Figures 2B-2D and S2).

Our analysis identified 53891 AS events across all genotypes, approximately 40% of which were alternative exons and a similar percentage was retained introns (Figures S2A and S2B). PSI (Ψ) values were used to quantify the mapped AS events in each sample (where [PSI] is the Percent Spliced In, i.e. the fraction of reads for an mRNA that includes the alternative sequence of the event).

Principal component analysis (PCA) of PSI values for alternative exon events demonstrated distinct grouping of genotypes carrying the *rpS12^G97D^*mutation, indicative of different AS patterns dependent on RpS12 function (PC1 exhibiting 19.95 % data variability) (Figure 2B). Furthermore, distinct grouping was observed between the *Rp*^+/−^ mutants and the remaining genotypes (PC2 13,69%), suggesting the presence of different AS events in *Rp*^+/−^ cells that depend on both RpS12 and Xrp1 function (Figure 2B). Note that also PCA based on the PSI values of all splicing events demonstrated distinct grouping of the genotypes carrying the *rpS12^G97D^* mutation (PC1 exhibiting 17.4 % data variability) (Figure S2C).

Analysis of PSI values for *Xrp1* splicing events revealed that exon 4 is highly included in the *Xpr1* mRNA across all samples analyzed (PSI values ranging from 92 to 99 percent) (Figures 2C, S2D and Supplementary Table 1).

Interestingly, *Rp*^+/−^ mutants exhibited a significant increase in exon 4 skipping, reducing PSI values compared to wild-type controls (Figures 2C and S2E). This skipping occurred independently of Xrp1, as the heterozygous *Xrp1^M273^*^/+^ null allele (*Xrp1*^+/−^) did not alter exon 4 PSI values in *RpS3*^+/−^ mutants (Figure 2C). Unexpectedly, the homozygous *rpS12^G97D^* mutation restored exon 4 inclusion in *RpS3*^+/−^ mutants to levels comparable to or exceeding wild-type PSI values, suggesting an RpS12-dependent mechanism of splicing regulation in *Rp*^+/−^ mutants (Figures 2C and S2E). Alternative exon 4 inclusion and skipping levels in the different genotypes are visualized using Sashimi plots, with numbered arcs quantifying splice junction reads (Figure 2D).

Semi-quantitative RT-PCR analysis, followed by gel electrophoresis, confirmed reduced exon 4 inclusion in both *Rp*^+/−^ mutants (*RpS3*^+/−^ and *RpS17*^+/−^) compared to wild-type cells (Figure 3A). The exon 4 skipping persisted in the presence of the heterozygous dominant *Xrp1^M273^*null mutation, demonstrating Xrp1 independence (Figure 3A). Furthermore, the *rpS12^G97D^*mutation increased exon 4 inclusion in *Rp*^+/−^ mutants, and notably also in wild-type cells (Figures 3A and S2E). This analysis suggests that RpS12, via exon 4 skipping, promotes the production of the RC transcript, encoding the Xrp1^short^ protein isoform (Figure 3B).

**Figure 3.**
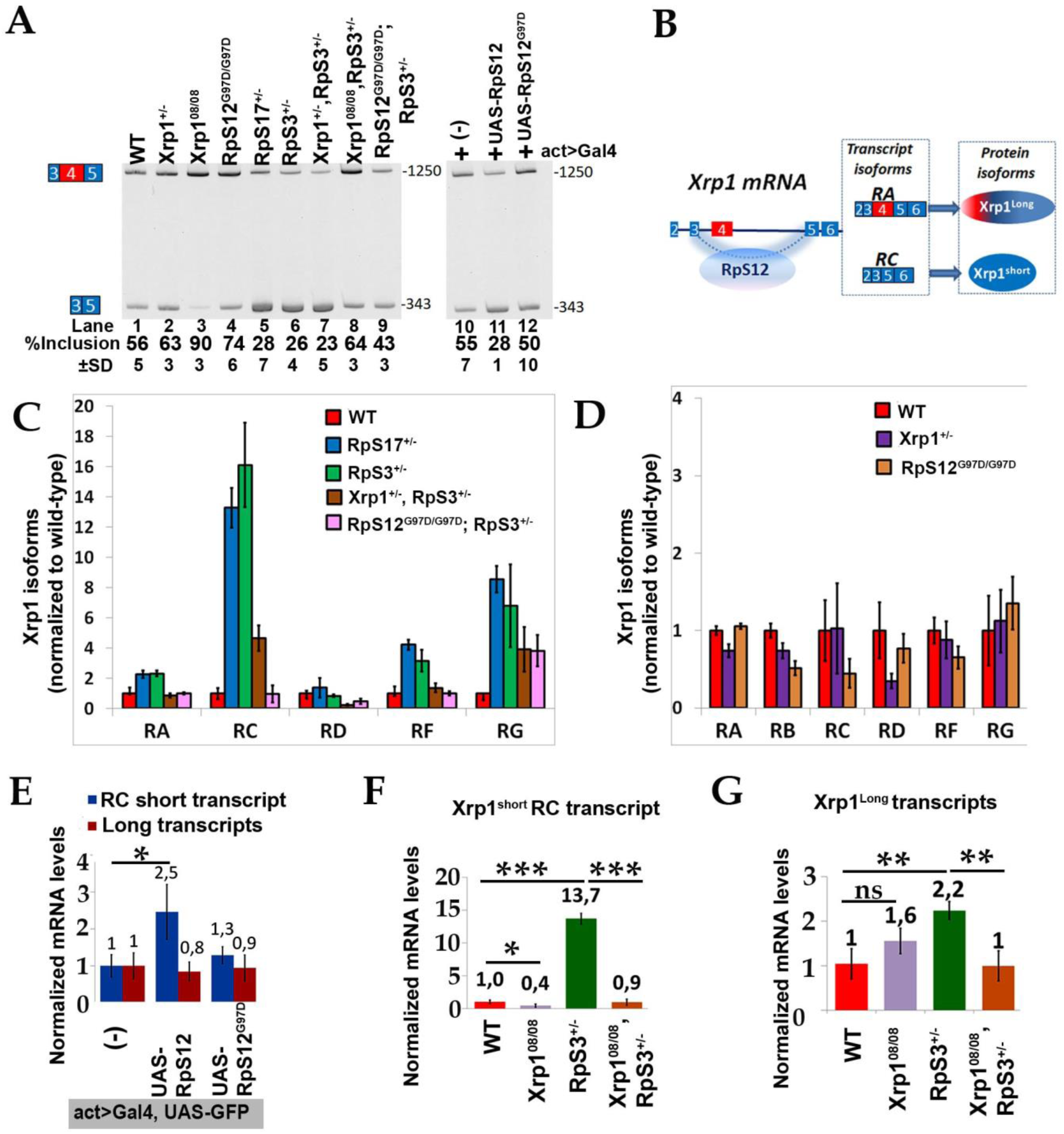
RpS12, through exon 4 skipping, induces expression of Xrp1^short^ isoform, which is required for the competitive elimination of the cells. (A) Analysis by semi-quantitative RT-PCR and gel electrophoresis of the splicing event of interest (Xrp1-Exon 4) across the different genotypes. Primers used anneal in exons 3 and 5, which simultaneously amplify all the isoforms that either include or exclude the exon 4. The % inclusion values represent the quantification of the inclusion band (1250bp) relative to the total signal from both the inclusion and skipping (343bp) bands. Error bars, based on the standard deviation (±SD) values from the corresponding biological replicates, are depicted below. Product lengths (bp) are marked on the right of each picture. (Genotypes shown per lane: *wild type* (1^st^ lane), *Xrp1^M273/+^* (2^nd^ lane), *Xrp1^08/08^* (3^rd^ lane), *RpS12^G97D/G97D^* (4^th^ lane), *RpS17^+/−^* (5^th^ lane), *RpS3^+/−^* (6^th^ lane), *Xrp1^M273/+^, RpS3^+/−^* (7^th^ lane), *Xrp1^08/08^,RpS3^+/−^*(8^th^ lane), *RpS12^G97D/G97D^, RpS3^+/−^* (9^th^ lane), *act>Gal4, UAS-GFP/+* (10^th^ lane), *act>Gal4, UAS-GFP/UAS-RpS12* (11^th^ lane), *act>Gal4, UAS-GFP/UAS-RpS12^G97D^* (12^th^ lane). (B) Model showing the RpS12-mediated splicing of Xrp1, which leads to exon 4 skipping and the production of the RC transcript, which encodes the Xrp1^short^ isoform. RA transcript encodes the long Xrp1 isoform. (C) Fold change of five of the Xrp1 mRNA isoforms normalized to wild-type levels. Graph depicts the ratio of transcripts per million of each isoform in indicated genotypes normalized to transcripts per million of each isoform in wild type cells. Error bars represent ±1 SD derived from RNAseq data. RE and RB isoforms are not included due to their low expression levels. Bar colors of genotypes: *wild-type* in red, *RpS17^+/−^* in blue, *RpS3^+/−^* in green, *Xrp1^+/−^, RpS3^+/−^*in brown and *RpS12^G97D^/^G97D^,RpS3^+/^*^-^ in pink. (D) Fold change of the Xrp1 mRNA isoforms normalized to wild-type levels. Graph depicts the ratio of transcripts per million of each isoform in indicated genotypes normalized to transcripts per million of each isoform in wild type cells. Error bars represent ±1 SD derived from RNAseq data. Bar colors of genotypes: *wild-type* in red, *Xrp1^+/−^*in purple and *RpS12^G97D/G97D^* in orange. (E) qRT-PCR measurements of the Xrp1 RC short transcript (blue columns) and of the Xrp1 long isoforms (red columns) from wing discs overexpressing with the act-Gal4 driver the reporter UAS-GFP, alone or together with UAS-RpS12 or UAS-RpS12^G97D^ transgenes. For the RC transcript the forward primer anneals at the junction of the exon 3-exon 5 and the reverse primer inside the exon 5. For the long transcripts the forward primer anneals at the junction of the exon 4-exon 5 and the reverse primer inside the exon 5. RpS12 overexpression upregulates two fold the RC transcript of the Xrp1, but not the long transcripts. The mutant RpS12^G97D^ is not sufficient to induce Xrp1. Error bars represent ±1 SD. p-values are calculated by Student’s t-test (* p< 0.05, ** p< 0.01, *** p< 0.001) (F) qRT-PCR measurements of the RC Xrp1^short^ isoform (using the same primer set as in e panel) in indicated genotypes. *Xrp1^08/08^* mutation reduces Xrp1^short^ transcript levels in wild-type cells and abrogates Xrp1^short^ upregulation in RpS3^+/−^ mutants. Error bars represent ±1 SD. p-values are calculated by Student’s t-test (* p< 0.05, ** p< 0.01, *** p< 0.001) (G) qRT-PCR measurements of the Xrp1 long isoforms (using the same primer set as in e panel) in indicated genotypes. *Xrp1^08/08^* mutation does not reduce Xrp1^long^ transcript levels in wild-type cells, but abrogates Xrp1^long^ upregulation in RpS3^+/−^ mutants. Error bars represent ±1 SD. p-values are calculated by Student’s t-test (* p< 0.05, ** p< 0.01, *** p< 0.001) See also Figure S3

Given that these results are based on the loss-of-function *rpS12^G97D^*allele, we investigated whether RpS12 overexpression is sufficient to induce exon 4 skipping, thus producing higher Xrp1 levels (Figure 1A). Semi-quantitative RT-PCR analysis showed that overexpression of wild-type RpS12 in wild-type cells replicated the exon 4 skipping phenotype observed in *Rp*^+/−^ mutants (Figure 3A, lanes 10-12). Similarly, qPCR analysis confirmed that RpS12 overexpression specifically upregulated the Xrp1^short^ isoform without affecting the Xrp1^long^ isoform (Figure 3E), suggesting that Xrp1^short^ may be sufficient to induce loser status. In contrast, overexpression of the RpS12^G97D^ mutant had no effect (Figures 3A and 3E), confirming the loss-of-function nature of the *rpS12^G97D^*allele. Therefore, both *loss-* and *gain-of-function* experiments support RpS12’s specialized role in regulating the alternative splicing of *Xrp1* exon 4 in both wild-type and *Rp*^+/−^ mutant cells.

The *Xrp1^08^* allele, previously shown to prevent the competitive elimination of Rp^+/−^ cells, has an intronic mutation near the 3’ of exon 4, disrupting a conserved putative splice enhancer[22] (Figure S3E). qPCR and semi-quantitative RT-PCR verified that *Xrp1^08^* homozygous cells exhibit specific reduction of Xrp1^short^ transcripts, without affecting long transcripts (Figures 3A, 3F and Suppl. Data in [22]). The reduced exon 4 inclusion in *RpS3*^+/−^ mutants is restored to wild-type levels in the presence of the *Xrp1^08/08^*allele (Figure 3A). Interestingly, in *RpS3*^+/−^ mutants the presence of the *Xrp1^08/08^* allele reduced the levels of both isoforms (Figures 3F and 3G). This result shows that the Xrp1^short^ isoform is involved in its transcriptional autoregulation in *Rp*^+/−^ cells, thus increasing both isoforms. Complementary, isoform analysis of Xrp1 transcripts using RNAseq data[21, 27] supported the AS analysis conclusions and revealed additional regulatory information. Transcriptional autoregulation in *Rp*^+/−^ cells by Xrp1 is evident for all transcript isoforms, with the RA long isoform being the most abundant in all genotypes (Figures 3C, and S3B-S3C). RA isoform displays two-fold upregulation in *Rp*^+/−^ mutants and its levels exhibit the same dependency on both Xrp1 and RpS12 function (Figures 3C, and S3B-S3C). Conversely, the less abundant RC short isoform, which is transcribed from the same promoter as RA, shows the highest induction in *Rp*^+/−^ cells. Expression of RC isoform is fully dependent on RpS12 function, but partially on Xrp1 autoregulation (Figures 3C, and S3B-S3C). This suggests that RC isoform in *Rp*^+/−^ cells depends on Xrp1 autoregulation, but in addition it is regulated by the RpS12-dependent mechanism. The RG transcript isoform is additionally regulated by an RpS12- and Xrp1-independent mechanism in *Rp*^+/−^ cells, implicating post-transcriptional control via its 3’UTR (Figures 3C, and S3B-S3C).

In wild-type cells, Xrp1 partially autoregulates its transcripts, with the RA (long) isoform showing modest feedback control (Figure 3D). However, the RC (short), RF and RG isoforms escape this regulation, maintaining their expression independent of Xrp1 activity (Figures 3D, and S3B-S3C). Notably, both *Xrp1^08^* and *rps12^G97D^* alleles selectively reduce the short isoform in wild-type cells, while in *Rp*^+/−^ cells the same alleles affect the levels of both isoforms (Figures 3D, and S3B-S3C). Given that Xrp1 levels are increased in *Rp*^+/−^ cells through a transcriptional autoregulatory mechanism[22, 24, 25], these data suggest that the Xrp1^short^ isoform, derived from RpS12-dependent splicing, is involved in autoregulation of the long isoform in *Rp*^+/−^ cells, but this is not happening in wild-type cells. This is consistent with RpS12 overexpression preferentially increasing short transcripts, but not long ones (Figure 3E).

Interestingly, while both AS and isoform analysis reveal that the RA transcript encoding the Xrp1^long^ isoform is the most abundant in *Rp*^+/−^ wing discs, only the short isoform is detected by Western blot (Figure S3D). Interestingly, all transcripts encoding the Xrp1^long^ isoform contain upstream open reading frames (uORFs)[48], which negatively affect the translational efficiency of the main ORF [35, 36]. Therefore, the RA isoform is under translational repression in both wild-type and *Rp*^+/−^ cells. In contrast, all the transcripts encoding the Xrp1^short^ isoform lack uORFs, and ribosomes can directly scan and efficiently initiate translation of the main ORF.

Altogether, our data support a model where the ribosomal protein RpS12 is involved in Xrp1 splicing regulation by promoting exon 4 skipping and producing the RC transcript. This event is required for generating the Xrp1^short^ protein isoform, which along with the Irbp18 protein, is involved in transcriptional autoregulation[24], necessary for *Rp*^+/−^ cells to adopt their loser identity (Figure S3A). Importantly, these data show that, while proteotoxic stress can activate Xrp1 and cell competition in other contexts, in *Rp*^+/−^ cells, RpS12 dependent production of the Xrp1^short^ isoform is the key driver of cell competition.

### Overexpression of either Xrp1 isoform is sufficient to transform wild-type cells into losers

While the role of Xrp1 in cell competition is well established across various contexts, whether Xrp1 is sufficient to induce loser status remained unclear[9]. Overexpression of either Xrp1 isoform can trigger part of the Rp^+/−^ responses and lead to massive autonomous cell death (Figures 4A, and S4A, S4B)[24, 31, 32, 38, 49]. Indeed, clonal overexpression of either the Xrp1^short^ or Xrp1^long^ isoform resulted in very few, small clones at the time of dissection (Figures 4B and S4C) [49]. Consequently, it was not possible to distinguish between autonomous and competitive cell death in these overexpression scenarios.

**Figure 4.**
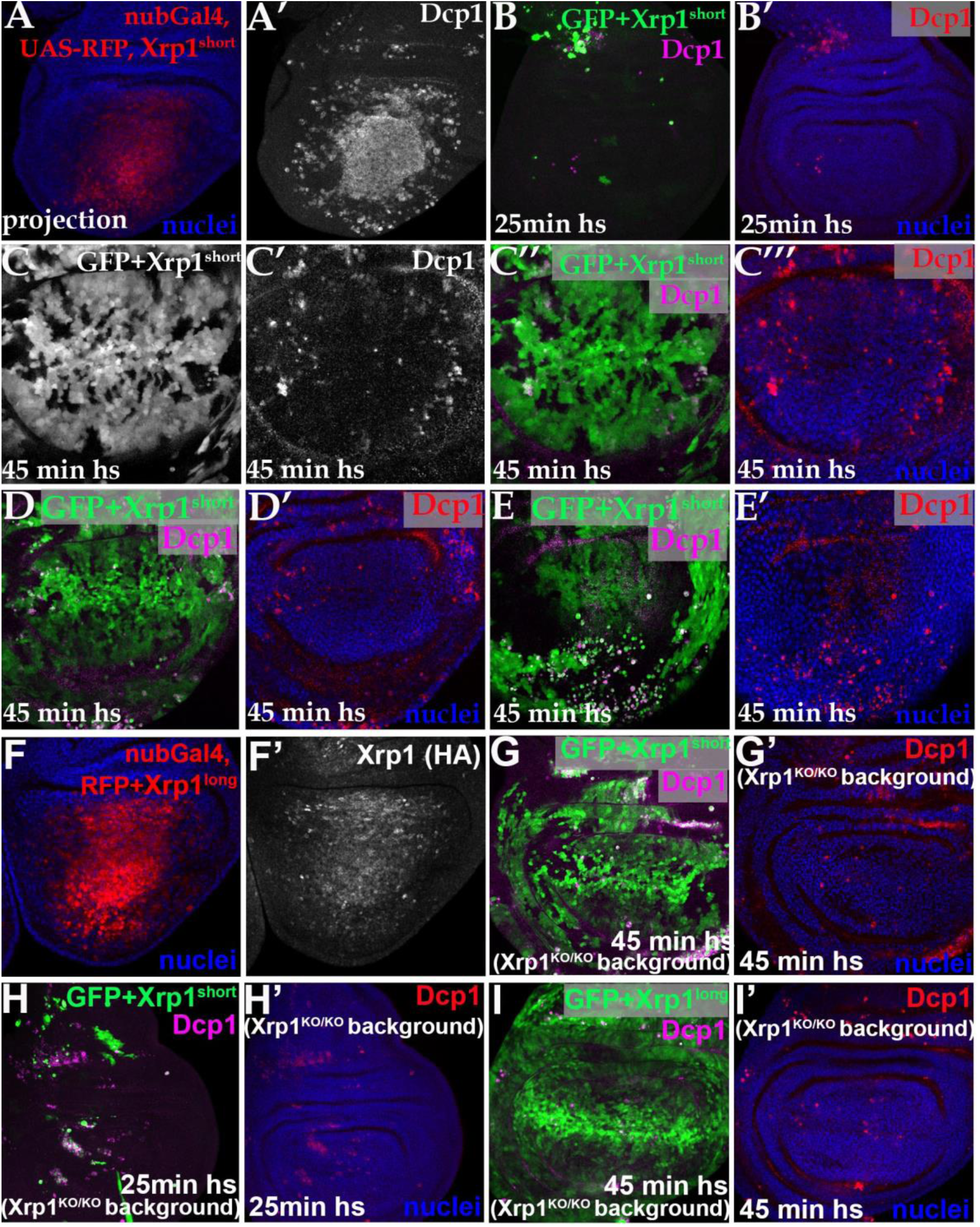
Overexpression of either Xrp1 isoform is sufficient to transform wild-type cells into losers. Confocal images of wing imaginal discs of the indicated genotypes. Panels (A), (B) and (H) show projections of active Dcp1 staining of the central disc-proper region. All the other panels show single confocal planes. (A) nub-Gal4 drives the co-expression of RFP (red, A) together with Xrp1^short^ in the pouch area of the wing. Xrp1^short^ overexpression resulted in massive cell death (Dcp1, A’). (B) Clones overexpressing Xrp1^short^ together with GFP (green, B) in wing discs did not survive (Dcp1, magenta in B and red in B’). Clones were induced by 25 min heat shock. (C) Upon longer heat-shock duration time (45 min), Xrp1^short^ overexpression in clones marked with GFP (C and green in C’’) in wild-type discs, led to the survival of the clones that exhibited boundary cell death (Dcp1 in C’, magenta in C’’ and red in C’’’). (D and E) Example of a wild-type disc where the disc proper area is covered by clones of cells (marked green with GFP, D) overexpressing Xrp1^short^, which present few boundary cell death (Dcp1, magenta in D and red in D’). In the peripodial membrane of the same disc, Xrp1 ^short^ over-expressing clones (green, E) present extensive boundary cell death. (Dcp1, magenta in E and red in E’). Clones were induced by 45 min heat shock. (F) nub-Gal4 drives the co-expression of RFP (red, F) together with Xrp1^short^ in the pouch wing area, which results in the induction of the endogenously HA-tagged Xrp1 allele (F’) (G) Clones of cells, marked with GFP (green, G), overexpressing the Xrp1^short^ isoform in a homozygous null Xrp1^KO^ background exhibit boundary cell death (Dcp1, magenta in G and red in G’). Clones induced by 45 min heat shock. (H) Xrp1^short^ overexpression in clones (GFP in green, H) in discs in a homozygous null Xrp1^KO^ background, led to massive cell death (Dcp1, magenta in H and red in H’). Clones induced by 25 min heat shock. (I) Example of a wild-type disc where the disc proper area is covered by clones of cells (marked with GFP, green in D) overexpressing Xrp1^long^, which present few boundary cell death (Dcp1, magenta in D and red in D’). See also Figure S4L, where in the peripodial membrane of the same disc, Xrp1^long^ over-expressing clones present extensive boundary cell death. Clones were induced by 45 min heat shock. See also Figure S4

Previous work has shown that merging loser clones facilitates their survival, allowing their recovery in mosaics with winner cells[50, 51]. Thus, we extended the heat shock duration to increase recombination frequency and generate larger areas of Xrp1^short^-overexpressing cells. We obtained many discs where the area of Xrp1^short^-overexpressing loser cells was large enough to observe cell death at their boundaries, as shown in a representative disc in Figure 4C. Consistent with active cell competition, the basal layer of the same discs contained numerous apoptotic loser cells resulting from loser extrusion by surrounding wild-type cells (Figure S4D). Furthermore, we retrieved many discs where Xrp1^short^ losers covered almost the entire disc proper and had a small number of apoptotic cells, preferentially located at the boundaries (Figures 4D and S4E). In contrast, in the peripodial layer (the thin squamous epithelium above the disc proper layer) of the same discs, we found an increased number of apoptotic losers adjacent to wild-type cells (Figures 4E and S4F). This difference highlights a key characteristic of cell competition, where competitive cell death of loser cells is more pronounced in a heterogeneous environment (e.g peripodial in Figure 4E) compared to a relatively homogeneous environment (disc proper in Figure 4D). Apoptotic loser cells on the same disc were extruded basally as expected since cell competition was still active in this disc (Figures S4G, control experiment in S4H). Notably, while ubiquitous RpS12 overexpression is viable, we never recovered flies with ubiquitous Xrp1 overexpression, suggesting that RpS12-overexpressing cells might activate an adaptive response that renders them resistant to Xrp1-induced cell death.

Xrp1^long^ overexpression has been shown to activate a transcriptional reporter of Xrp1 [22, 24]. Using the endogenously HA-tagged allele of Xrp1, we found that overexpression of either Xrp1^short^ or Xrp1^long^ isoform upregulated Xrp1 (Figures 4F, S4I, control disc in S4J). Note that both UAS transgenes, include only the coding regions of the Xrp1 isoforms and do not include 5’ and 3’ untranslated regions (UTRs)[52]. Based on these findings, we overexpressed the Xrp1^short^ isoform in the absence of the long isoform to determine whether Xrp1^short^ is sufficient to trigger cell competition. Indeed, applying prolonged heat-shock, Xrp1^short^ overexpression in a homozygous null Xrp1^KO^ background resulted in discs where almost the entire disc proper was covered by loser cells, which also exhibited boundary cell death (Figures 4G and S4K). Still, with a shorter heat shock duration, the Xrp1^short^ overexpressing clones did not survive (Figure 4H).

Finally, we also investigated whether the Xrp1^long^ isoform is sufficient to induce competition by overexpressing it in a homozygous Xrp1^KO^ background. Similar to Xrp1^short^, we only retrieved clones after a prolonged heat-shock duration. In many of these discs Xrp1^long^ losers covered almost the entire disc proper and had a small number of apoptotic cells, preferentially located at the boundaries (Figure 4I). In contrast, in the peripodial layer of the same discs, an increased number of apoptotic losers were found adjacent to wild-type cells (Figure S4L).

### Xrp1^long^ isoform is not required for *Rp*^+/−^ cell competition

To examine the role of the Xrp1^long^ exons in *Rp*^+/−^ dependent cell competition, we used the previously generated *Xrp1^Exlong^*allele, where the long exons are deleted (Figure S3E)[52]. The *Xrp1^Exlong^* allele expresses the Xrp1^short^ isoform from the Xrp1-RD transcript [52]. Consistent with the above findings, *RpS17*^+/−^ cells in mosaics with wild-type cells still exhibited boundary cell death in the presence of the homozygous *Xrp1^Exlong^*allele (Figure 5A). Notably, *Rp*^-/-^ cells were observed in this context (Figure S5A), which had been previously observed in homozygous *Xrp1*^null^ background or upon apoptosis inhibition [21, 53], supporting the distinct requirements for *Rp*^-/-^ elimination versus competitive cell death [53].

**Figure 5.**
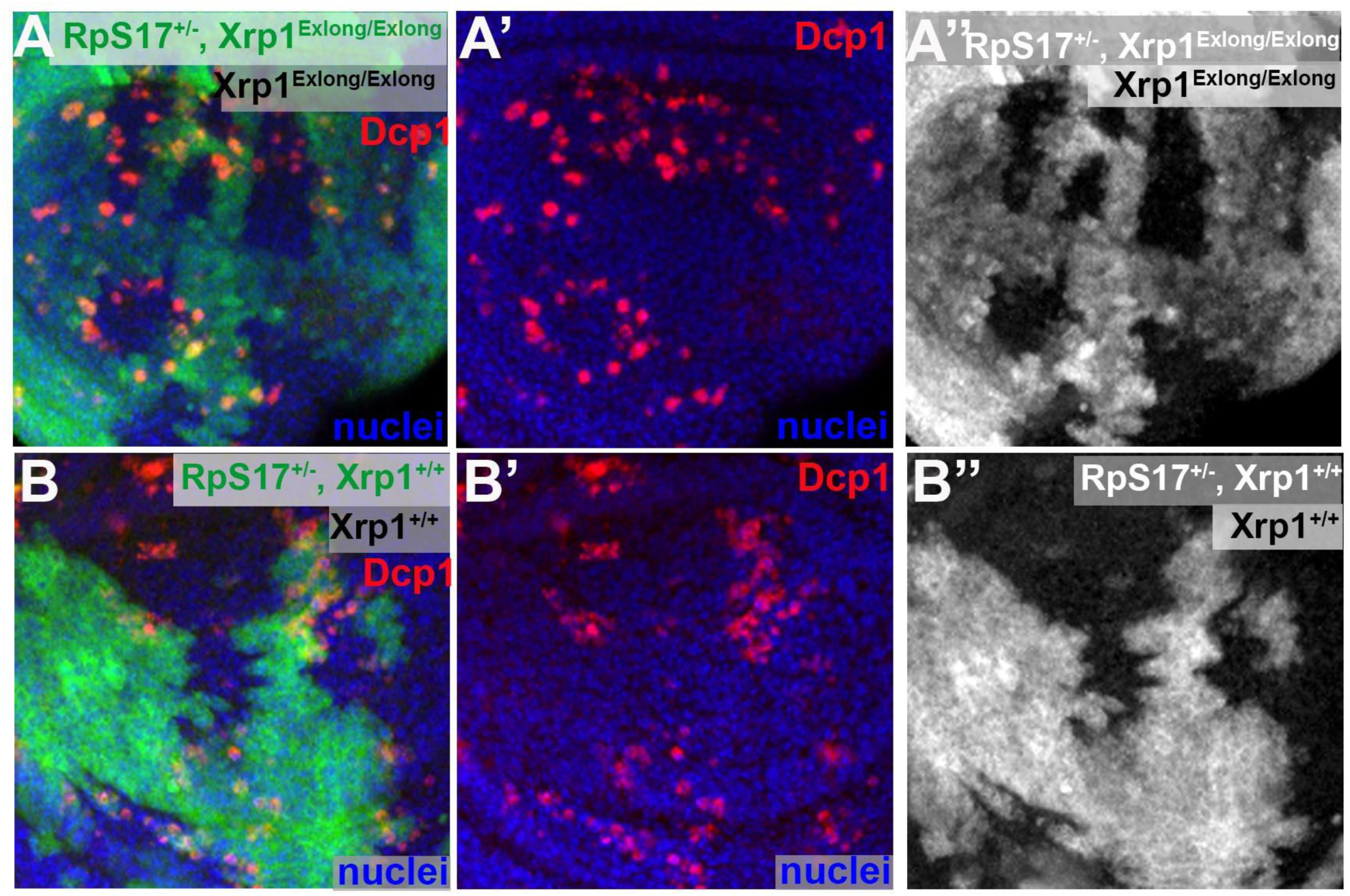
Xrp1^short^ isoform is sufficient to induce cell competition in *Rp*^+/−^ cells. Confocal images of wing imaginal discs of the indicated genotypes. All panels show single confocal planes. (A) Cell competition still occurs between *RpS17*^+/−^ cells and wild-type cells in wing disc homozygous for the Xrp1^Exlong^ allele. In Xrp1^Exlong^ allele the two thirds of the Xrp1 gene is deleted, predicted to abolish expression of the Xrp1^long^ isoform. In Xrp1^Exlong^ flies some residual Xrp1 transcript is detected (∼8% of WT levels)[52], possibly reflecting expression of the Xrp1 short isoform. (B) Control experiment presenting cell competition between *RpS17*^+/−^ cells and wild-type cells in wing disc with intact the Xrp1 locus. See also Figure S5

Interestingly, *RpS17*^+/−^,Xrp1^Exlong/Exlong^ cells occupied a larger area in mosaics with wild-type cells compared to *RpS17*^+/−^ (Figures 5A and 5B). While more rigorous quantification is needed, this observation suggests that cell competition persists in the absence of the long Xrp1 exons, while the growth of the *Rp*^+/−^ cells is restored. These results suggest that the Xrp1^short^ protein isoform is both necessary and sufficient for driving “loser” identity in both *Rp*^+/−^ and wild-type cells. In addition, while Xrp1^long^ isoform overexpression can induce the loser status of wild-type cells, this isoform is dispensable for *Rp*^+/−^ cell competition.

### RNA-binding protein Syncrip is involved in Xrp1 regulation

To gain insights into the mechanism underlying Xrp1 alternative splicing, we performed a proteomic analysis to identify changes in RNA-binding proteins (RBP) in *Rp*^+/−^ cells. The equilibrium of RBPs in cells is finely regulated, and alterations in their expression levels can influence RNA processing, including splicing[54]. Therefore, we hypothesized that changes in RBP levels involved in Xrp1 splicing would occur independently of Xrp1 itself. To test this, we compared protein expression profiles in wing discs from: a) *wild*-*type*; b) *RpS3*^+/−^; and c) *Xrp1*^+/−^, *RpS3*^+/−^.

Hierarchical clustering of differentially expressed proteins identified three distinct clusters with unique functional enrichments (Figures 6A and 6B). Cluster I comprised 232 proteins downregulated in *RpS3*^+/−^ cells compared to wild-type, independent of Xrp1. This cluster was enriched for Gene Ontology terms associated with cytoplasmic translation, axon guidance, septate junction assembly, and dendrite self-avoidance. Cluster II included 134 proteins upregulated in *RpS3*^+/−^ cells relative to wild-type, also independent of Xrp1. This cluster was enriched for terms related to ribosomal RNA processing and ribosome biogenesis, confirming previous findings showing that ribosome biogenesis intermediates accumulate independently of Xrp1 in *Rp*^+/−^ cells[21, 25]. These proteins in Clusters I and II represent potential candidates contributing to RpS12-dependent splicing of Xrp1.

**Figure 6.**
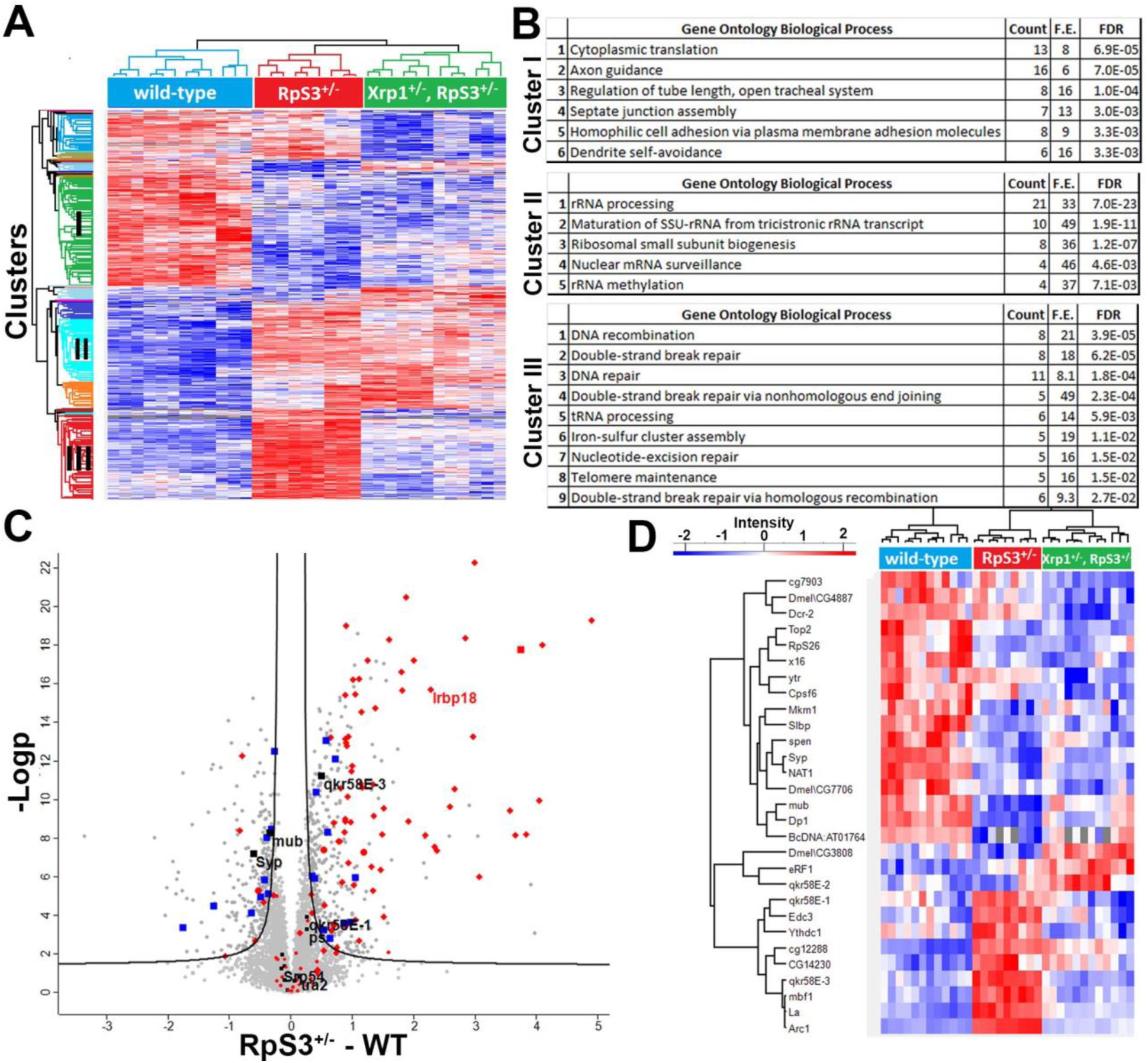
Proteomic analysis reveals Xrp1-dependent and independent changes of *Rp*^+/−^ cells, including alterations in RNA-binding proteins. (A) Hierarchical cluster analysis of differential expressed proteins between *wild-type*, *RpS3*^+/−^ and *Xrp1^+/−^, RpS3*^+/−^ samples. Blue indicates gradually downregulated proteins and red gradually upregulated proteins. Three selected clusters presenting distinct behavior with respect to proteomic changes are highlighted in the dendrogram. Cluster I: Xrp1-independent downregulated proteins in *Rp*^+/−^ cells, Cluster II: Xrp1-independent upregulated proteins in *Rp*^+/−^ cells; Cluster III: Xrp1-dependent upregulated proteins in *Rp*^+/−^ cells, (B) Functional analysis of protein annotation terms resulted in multiple categories of biological processes that were enriched in the three clusters. The enriched terms and the corresponding enrichment factor and *p*-value are shown. (C) Volcano plot showing results of differential expression analysis comparing proteins from wild-type wing discs and *RpS3*^+/−^ mutant wing discs. The vertical line differentiates the upregulated and the downregulated proteins. Square blue symbols indicate mRNA-binding proteins differentially expressed in *RpS3*^+/−^ cells. Square black symbols indicate RBP previously shown to interact with Xrp1 transcript [46]. Diamond red symbols indicate differentially expressed proteins in *Rp*^+/−^ cells, whose mRNA was also differentially expressed in *Rp*^+/−^ cells[21]. Almost all of them belong to Cluster III, Irbp18 is one of them. Note that less than 250 genes were differentially expressed based on our previous RNAseq analysis[21]. (FDR=0.01) (D) Heatmap showing significantly altered RNA-binding proteins between *wild-type* wing discs, *RpS3^+/−^* discs and *Xrp1^+/−^, RpS3*^+/−^ discs. Blue indicates gradually downregulated proteins and red gradually upregulated proteins.

Finally, cluster III comprised 180 proteins upregulated in RpS3^+/−^ cells in an Xrp1-dependent manner. This cluster showed enrichment for pathways involved in DNA recombination, DNA repair, DNA damage, and iron-sulfur cluster assembly. This confirms previous RNA-seq data showing that Xrp1 transcriptionally activates these pathways in *Rp*^+/−^ cells[21]. The presence of Irbp18 in Cluster III, further validates our proteomic analysis (Figure 6C).

Our proteomic analysis identified numerous differentially expressed RNA-binding proteins (RBPs) across the different genotypes (Figures 6C and 6D). Notably, Syncrip (Syp), an RBP previously shown to bind Xrp1 mRNA in S2 cells[46], was downregulated in *RpS3*^+/−^ cells, and this downregulation persisted in *Xrp1*^+/−^, *RpS3*^+/−^ cells (Figures 6C and 6D). To investigate Syp’s role in Xrp1 regulation in *Rp*^+/−^ cells, we initially examined whether reducing Syp levels in wild-type cells was sufficient to induce Xrp1 expression. Silencing Syp in the posterior compartment of wild-type wing discs using two different RNAi lines resulted in Xrp1 induction with both lines (Figures 7A, C, and S6A), suggesting that Syp negatively regulates Xrp1. However, the strength of Xrp1 induction varied between the RNAi lines, with one line showing a more pronounced effect (Figures 7A and 7C). Importantly, in RpS18^+/−^ discs, only the stronger RNAi line led to a further increase in Xrp1 levels (Figures 7B, and S6B). This observation aligns with the already reduced Syp levels in *Rp*^+/−^ cells, which render these cells less responsive to Syp-RNAi expression, and thus allowing only the strong RNAi line to increase Xrp1 levels in those cells.

**Figure 7.**
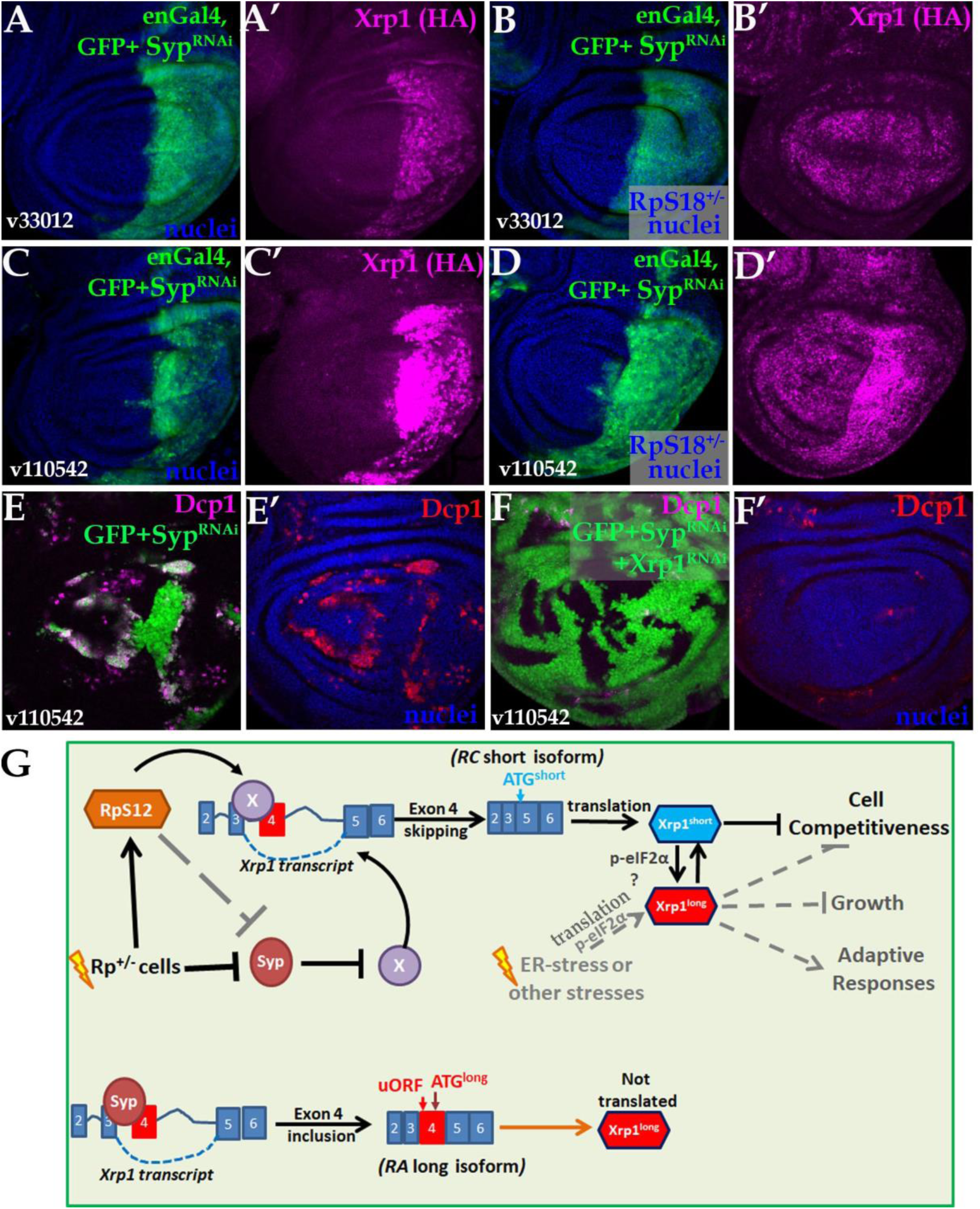
RNA-binding protein Syncrip is involved in the regulation of Xrp1 and cell competition. Confocal images of wing imaginal discs of the indicated genotypes. Panels (E) and (F) show projections of Dcp1 staining of the central disc-proper region. All the other panels show single confocal planes. (A) en-Gal4 drives the co-expression of GFP (A) together with Syp-RNAi (v33012 line) in the posterior domain of a wild-type wing disc, resulting in the induction of the endogenously HA-tagged Xrp1 allele (A’) (B) en-Gal4 drives the co-expression of GFP (B) together with Syp-RNAi (v33012 line) in the posterior domain of a *RpS18*^+/−^ wing disc, which does not show any further induction of the endogenously HA-tagged Xrp1 allele (B’) (C) en-Gal4 drives the co-expression of GFP reporter (C) together with Syp-RNAi (v110542 line) in the posterior domain of a wild-type wing disc, which results in the strong induction of the endogenously HA-tagged Xrp1 allele (C’) (D) en-Gal4 drives the co-expression of GFP (D) together with Syp-RNAi (v110542 line) in the posterior domain of a *RpS18*^+/−^ wing disc, which results in the strong induction of the endogenously HA-tagged Xrp1 allele (D’) (E) Clonal Syp depletion by RNAi (v110542) (green in B) in wild-type discs led to boundary cell death (Dcp1, magenta in E and red in F). Clones were induced by 45 min of heat-shock. No clones were retrieved upon 25 minutes heat-shock (not shown). (F) Boundary cell death (Dcp1, magenta in F and red in F’) in clones overexpressing the Syp-RNAi (v110542) was completely abolished upon depletion of Xrp1 by RNAi (green in F). (G) Proposed model: In *Rp*^+/−^ cells, RpS12 promotes exon 4 skipping of the *Xrp1* mRNA, producing the RC transcript. This transcript lacks uORF in its 5’ UTR and is efficiently translated, producing the Xrp1^short^ isoform. Xrp1^short^ isoform is necessary and sufficient to transform cells into loser and induce cell competition in both *wild-type* and *Rp*^+/−^ cells. Xrp1^short^ protein increases the levels of all transcripts, including the *Xrp1^long^* transcript. RA isoform encoding the Xrp1^long^ isoform is the most abundant transcript in both wild-type cells and *Rp*^+/−^ cells, but fails to produce detectable protein, probably due to uORFs within its 5’ untranslated region. In *Rp*^+/−^ cells, Xrp1^short^ could induce Xrp1^long^ translation due to p-eIF2α induction. Therefore, Xrp1 ^long^ isoform may be responsible for the eIF2α-dependent adaptive responses in *Rp*^+/−^ cells and for their reduced growth. Other stresses (such as ER stress), could induce Xrp1^long^ isoform translation through uORF-dependent mechanisms and subsequently increase the short isoform through autoregulation. We propose that Syncrip may inhibit exon 4 skipping, by antagonizing the binding of other RBP on Xrp1. In *Rp*^+/−^ cells, Syp levels are reduced through an RpS12-dependent mechanism, thus allowing exon 4 skipping by another RBP. Syp could also inhibit Xrp1 expression by a splicing-independent and/or RpS12-independent mechanism. See also Figure S6 and S7

Consistent with Xrp1 expression, Syp knockdown conferred loser status on wild-type cells, inducing their death at the clone boundaries with wild-type cells (Figure S6C). The stronger Syp-RNAi line more effectively transformed cells into ‘losers’ during competition, with loser clones surviving only after prolonged heat shock (Figure 7E). Strikingly, co-depletion of Xrp1 completely abolished the competitive cell death induced by Syp depletion (Figure 7F). These results demonstrate that Syp levels, by regulating Xrp1 expression, mediate fitness differences in wild-type cells and likely contribute to the competitive disadvantage of *Rp*^+/−^ cells.

## DISCUSSION

### RpS12 dependent splicing of Xrp1, rather than proteotoxic stress, drives the *Rp*^+/−^ phenotypes

Similar to *Minute* syndrome in flies, heterozygous mutations in *Rp* genes in humans can lead to Diamond Blackfan Anemia, a ribosomopathy[1, 3]. Ribosomopathies arise from defects in ribosome synthesis or function and are typically characterized by hematopoietic and developmental abnormalities, along with a paradoxical increase in cancer risk[1, 3, 55–58]. Two key pathophysiological mechanisms are implicated in these diseases: p53 activation due to disrupted ribosome biogenesis, and translational alterations caused by reduced ribosome numbers or impaired ribosome function[3, 55, 59–61].

Interestingly, in *Drosophila*, Xrp1 is the major transcriptional target of p53 following irradiation with a crucial role in the DNA damage response and genome stability[62, 63]. However, p53 is not required for Xrp1 expression and cell competition in *Rp*^+/−^ cells [21, 53]. Therefore, in *Rp^+/−^* cells, Xrp1 appears to function analogously to mammalian p53[64]. The prevailing model of p53 activation in ribosomopathies posits that unassembled RPs interact with MDM2, inhibiting its ability to degrade p53[65]. *Rp^+/−^* cells in *Drosophila* also display an accumulation of ribosomal intermediates[25]. Thus, we previously proposed that the function of the mammalian Rp/MDM2/p53 signaling pathway is performed in *Drosophila* by an RpS12-dependent intermediate, or RpS12 itself, which activates Xrp1[64].

Here, we show that in *Rp*^+/−^ cells, RpS12 promotes the production of the Xrp1^short^ isoform through exon 4 skipping. The *rpS12^G97D^* mutant lacks this function. Remarkably, we found that RpS12 overexpression is sufficient to induce exon 4 skipping and Xrp1 expression, transforming wild-type cells into “losers”. In contrast, overexpression of RpS12^G97D^, RpS17, RpL14 or RpL13 proteins does not promote Xrp1 expression or cell competition, suggesting that RpS12 possesses a specialized signaling function. Therefore, *Minute* phenotypes arise not from the loss of ribosomal function, but from the gain of specific RpS12 activity.

Interestingly, DNA damage in human cells triggers a signaling pathway where SMG1 protein is released from TP53 mRNA. This allows RpL26 to bind p53 mRNA and recruit splicing factor SRSF7, promoting its alternative splicing [66]. This leads to the production of the shorter p53β isoform, which promotes cellular senescence[66]. In yeast, RpS12 has been implicated in senescence regulation[67]. While yeast lack homolog of p53, this highlights an evolutionary conserved response, where RPs regulate senescence through genome stability surveillance. The potential involvement of this Rp-dependent splicing mechanism in p53 regulation and senescence induction in human ribosomopathies remains unexplored.

Further research is needed to elucidate whether RpS12 acts directly as a *bona fide* splicing factor, like some of its homologs[68], or indirectly affects splicing, e.g. by translational regulation of a splicing factor (e.g Syp)[29, 69–74]. Interestingly, RpL10Ab homologs have been shown to regulate alternative splicing and participate in specialized translation of mRNAs[75, 76]. It will be interesting to investigate in the future the relation among RpL10A, RpS12 and Xrp1 alternative splicing during cell competition.

### The Xrp1^short^ isoform is necessary to induce the loser status

The role of the Xrp1^short^ isoform in cell competition is further supported by the *Xrp1^08^* allele, which reduces production of the short transcripts and prevents the competitive elimination of *Rp*^+/−^ cells. Our data support a model in which RpS12 produces the Xrp1^short^ isoform in *Rp*^+/−^ cells, which in turn, through autoregulation, increases levels of all Xrp1 transcripts. While the RA transcript, encoding the Xrp1^long^ isoform, is the most abundant in both wild-type cells and *Rp*^+/−^ cells, it fails to produce detectable protein, probably due to uORFs within its 5’ untranslated region (5’ UTR) [35, 36, 48] (Figures S3C and S3D). Conversely, all short transcripts, expressed at lower levels, lack uORFs and are readily translated by ribosomes upon induction in *Rp*^+/−^ cells. This may also explain why cells maintain low levels of short transcripts (Figure S3C). While 5’ UTR contributes to mRNA stability[77], PCA plots show that Rps12 regulates the splicing of multiple genes, in support of the notion that RpS12 affects the splicing of Xrp1 transcripts.

Xrp1 is activated by various other stresses, including increased p-eIF2α levels, and induces cell competition [9], while RpS12 is not always involved in Xrp1 induction [33]. Therefore, it will be crucial to investigate whether, under these stress conditions, Xrp1^long^ isoform translation is activated through uORF-dependent mechanisms, which could subsequently activate also the short isoform through transcriptional autoregulation.

As previously shown, *Rp*^+/−^ cells exhibit increased p-eIF2a levels due to Xrp1 induction [25, 31, 32, 40, 41]. Xrp1^short^ is the predominant protein isoform detected in *Rp*^+/−^ cells (Figure S3D), and p-eIF2a depletion does not affect its levels [25], suggesting that Xrp1^short^ isoform is responsible for increased PERK activity and p-eIF2a levels (Figure 7G). Indeed, overexpression of the Xrp1^short^ isoform can activate PERK/p-eIF2α response in wild-type cells [32]. While increased p-eIF2a levels generally suppress global protein synthesis, certain mRNAs containing specific uORF can be preferentially translated[48]. Previous results suggested a protective role for p-eIF2a in *Rp*^+/−^ cells[25, 40, 41]. Consequently, we endorse the idea that, in *Rp*^+/−^ cells, increased p-eIF2a levels promote the translation of the Xrp1^long^ isoform, which has a protective role, even if remains undetected by western blot analysis. Supporting this, Boutros et al. observed that, in *Drosophila* cell lines, targeting both isoforms via RNAi resulted in no noticeable phenotype [Supplementary data in [78]], as expected given that *Xrp1* null mutants are viable. Unexpectedly, depleting only the long Xrp1 transcript in these cell lines caused growth and viability defects [Figure 3D in [78]]. This suggests that, also in these cell lines, the Xrp1^long^ isoform has a protective role, possibly by mitigating the detrimental effects of the short isoform.

Future studies should investigate whether the reduced growth of *Rp*^+/−^ cells requires the Xrp1^long^ isoform, even though cell competition can still proceed without it. This will allow us to dissect the distinct contributions of the Xrp1 isoforms to the growth defects and competitive behavior of *Rp*^+/−^ cells, and possibly mechanistically separate the growth defect from reduced competitiveness.

### Xrp1 is sufficient to induce the loser status

Our study further demonstrates, for the first time, that Xrp1 is not only necessary but also sufficient to induce “loser” status in wild-type cells. This finding strongly supports the hypothesis that Xrp1 is the primary instigator of cell competition in *Rp*^+/−^ cells, rather than simply a component of their competitive cell death. Specifically, while both isoforms are sufficient to promote the loser identity in wild type cells when overexpressed, the short isoform is the one that is required in *Rp*^+/−^ cells. The Xrp1^long^ isoform may have role in other Xrp1-dependent cell competition contexts, where RpS12 is not involved[33].

Our proteomic analysis of *Rp*^+/−^ cells, coupled with the Xrp1 dependency assessment, provides a valuable resource for dissecting the complex effects of *Rp* haploinsufficiency. This approach enables us to: 1) distinguish between pathways directly affected by *Rp* mutations and those activated by Xrp1; 2) identify molecular components involved in RpS12-dependent Xrp1 splicing, such as Syncrip; and 3) uncover downstream effectors of Xrp1 that mediate cell competition and regulate growth. Therefore, this proteomic analysis will be a valuable dataset and will offer novel mechanistic insights into the molecular mechanisms underlying cell competition and the broader responses to ribosomal protein haploinsufficiency.

### RNA-binding protein Syncrip is involved in the regulation of Xrp1 and cell competition

Our proteomic analysis revealed decreased Syncrip (CG17838/Syp) levels in Rp^+/−^ cells, independent of Xrp1 (Figure 6), suggesting that Syp might regulate Xrp1 alternative splicing.

Syp, the Drosophila homolog of mammalian SYNCRIP/hnRNP Q, plays critical roles in post-transcriptional regulation, particularly in RNA processing, transcript stability, and localized translation [79–81]. Critically, we found that reducing Syp levels in otherwise wild-type cells was sufficient to trigger Xrp1-dependent cell competition (Figure 7). Notably, human Syncrip interacts with many RPs, including RpS12,[82] and with the spliceosome[83].

Previous studies have shown that Syp binds to the Xrp1 mRNA intron upstream of exon 4, along with several other RBPs, including mub, ps, qkr58E-1, tra2, and Srp54 (Figure S7D)[46]. Our proteomic analysis revealed increased mub levels in *Rp*^+/−^ cells (Cluster I), while the increased levels of qkr58E-1 in *Rp*^+/−^ cells depends on Xrp1 (Cluster III). These findings lead the intriguing suggestion that Syp regulates Xrp1 splicing by antagonizing the binding of the other RBPs (Figures S7A and S7C). In *Rp*^+/−^ cells, reduced Syp levels could permit increased binding of these other RBPs, leading to enhanced Xrp1 expression via exon 4 skipping (Figure S7B).

Future studies will investigate whether RpS12 directly or indirectly is involved in Syp downregulation in *Rp*^+/−^ cells, and whether Syncrip regulates Xrp1 through splicing or another post-transcriptional mechanism (Figure 7G). Interestingly, reduced Syncrip levels in haematopoietic stem cells (HSC) impair their competitive fitness when transplanted alongside wild-type HSCs [84]. Notably, Syncrip-depleted HSC exhibit increased levels of the CHOP transcription factor [84], the functional homolog of Xrp1 in mammals [9, 22, 24, 32, 36]. This suggests a conserved evolutionary role for Syncrip, and possibly for CHOP, in determining cellular adaptation and fitness.

Xrp1 has previously been shown to mediate the toxicity of another *Drosophila* RNA-binding protein, Cabeza[52, 85]. This raises the intriguing possibility that Xrp1, by virtue of its large mRNA size and RBP affinity[46], acts as a molecular sensor for the availability of multiple RBP (an “RBPostasis sensor”), promoting the elimination of cells with dysregulated RBP levels. Consistent with this hypothesis, Xrp1 drives stress and growth responses in cells with compromised spliceosome function[34].

In conclusion, we demonstrate that the activities of RpS12 and Xrp1, observed in *Rp*^+/−^ cells, are by themselves sufficient to transform cells into losers. We outline the underlying mechanism, revealing that RpS12 induces the short Xrp1 isoform production by alternative splicing regulation. The RpS12-dependent induction of the Xrp1^short^ isoform is essential for triggering cell competition in both *Rp*^+/−^ and wild-type cells. Collectively, our results establish a direct pathway linking *Rp*^+/−^ genotypes to Xrp1 expression. We show, for the first time, that Xrp1 expression, either the long or the short isoform, is sufficient to initiate cell competition in otherwise wild-type cells. Therefore, investigating the Xrp1-mediated responses driving *Rp*^+/−^ elimination in mosaics with wild-type cells can illuminate the mechanisms underlying the universal phenomenon of cell competition. Finally, understanding the mechanism of RpS12-dependent splicing in haploinsufficiency-associated phenotypes in *Drosophila*, may provide insights into two unexpected hallmarks of ribosomopathies: their tissue-specific defects and their increased cancer risk.

## METHODS

### RESOURCE AVAILABILITY

#### Lead Contact

Further information and requests for resources and reagents should be directed to and will be fulfilled by the Lead Contact, Marianthi Kiparaki.

#### Materials Availability

Most materials are commercially available. Further unique reagents generated in this paper are available from the lead contact without restriction.

#### Data and Code availability

The data supporting the findings of this study are available within the paper and its Supplementary Information. The mass spectrometry dataset will be available upon publication.

This paper does not report original code.

### EXPERIMENTAL MODEL

Species: Drosophila melanogaster.

Drosophila strains were generally maintained at 25°C on standard wheat flour-sugar food supplemented with soy flour and CaCl2, at 25°C with 50%–70% relative humidity on a 12-hour light/dark cycle. Sex of larvae dissected for most imaginal disc studies was not differentiated.

Full genotypes for all the experiments are listed in Supplementary Table 2. The following genetic strains were used: UAS-RpL13A.Flag (Bloomington 83684), UAS-RpS17 [86](FlyORF, F001268), UAS-GFP-RpL10Ab (Bloomington 42683), act>stop>Gal4, UAS-CD8-GFP/Cyo (gift from Christos Delidakis, UAS-SypRNAi (v110542 and v33012), UAS-Xrp1^short^ [52], UAS-Xrp1^long^ [52], Xrp1^KO^ [52], Xrp1^Exlong^ [52]. Other stocks are described in [21, 24, 25, 28]. Xrp1 protein expression was examined making use of an allele tagged with HA at the endogenous Xrp1 locus[24].

### METHOD DETAILS

#### Immunohistochemistry and Antibody Labeling

For antibody labeling, imaginal discs from late 3rd instar larvae were dissected in 1xPBS buffer and fixed in 4% formaldehyde in 1 x PEM buffer (1x PEM:100 mM Pipes, 1 mM EGTA, 1 mM MgCl2, pH 6.9). Fixed imaginal discs were 3x washed in PT (0.2% Triton X-100, 1xPBS) and blocked for at least 1hr in PBT buffer (0.2% Triton X-100, 0.5% BSA, 1 x PBS). Discs were incubated in primary antibody in PBT overnight at 4°C, washed three times with PT for at least 10 min each. The samples were incubated in secondary antibody in PBT overnight at 4°C (or for 3–4 hr at room temperature), and washed three times with PT for 5–10 min. After washes, samples were rinsed in 1 x PBS and the samples were incubated with the NuclearMask reagent (Thermofisher, H10325) for 10–15 min at room temperature. After washing 2 x with 1x PBS the imaginal discs were mounted in VECTASHIELD antifade mounting medium (Vector Laboratories, H-1000).

We used the following antibodies for staining: mouse anti-b-Galactosidase (J1e7, DSHB), rabbit anti-active-Dcp1 (Cell Signaling Techonology Cat#9578, 1:100), anti-HA Tag mouse monoclonal (Cell Signalling Cat#2367). Secondary Antibodies were Alexa Fluor conjugates (Thermofisher).

#### Clonal analysis

Genetic mosaics were generated using the FLP/FRT system employing inducible heat-shock FLP (hsFLP) transgenic strains [87, 88]. For making clones through mitotic recombination using inducible heat-shock FLP (hsFLP), *Rp*^+/−^ larvae were subjected to 60 min heat shock at 37 °C, 60 ± 12 hours after egg laying (AEL) and dissected 68-72 hr later. For making clones by excision of a FRT cassette, larvae were subjected to 20–60 min heat shock at 37 °C, 40 ± 12 AEL and dissected 68-72 hr later. Full genotypes for all figures and heat shock times are listed in Supplementary Table 2.

#### Image Acquisition and Processing

Confocal images were recorded using Leica Sp8 and Zeiss confocal microscopes using 20x and 40x objectives. Images were processed using Leica’s and Zeiss software, and Adobe Photoshop CS5 Extended.

#### Alternative splicing analysis

Splicing analysis of the RNA-Seq data (GEO accession numbers: GSE112864 and GSE124924) was performed using the Alternative Splicing and Transcript Tools (VAST-TOOLS) v2.2.2 (https://github.com/vastgroup/vast-tools), using Drosophila genome release dm6[89]. For all events, a minimum read coverage of 10 actual reads per sample was required, as described[90]. PSI values for single replicates were quantified for all types of alternative events, including single and complex exon skipping events (S, C1, C2, C3, ANN), alternative donor and acceptor splice choices (Alt5 and Alt3, respectively) and retained introns (IR-S, IR-C). A minimum dPSI of 5% was required to define differentially spliced events upon each mutant, along with a *p-value* < 0.05 calculated using 2-tailed unpaired Wilcoxon Mann–Whitney test. Sashimi graph was generated using IGV and Aligned BAM files (dm6 genome).

#### Salmon isoform quantification analysis

The same RNA-seq data was quantified using the Salmon software (v0.14.1)[91]. Transcript abundance was estimated using quasi-mapping-based mode. The analysis was performed using the pre-built transcriptome index generated from the Drosophila melanogaster genome (BDGP6.46 from Ensembl release 112), while setting the minimum score fraction parameter to 0.95 to retain only high-quality mappings. The output TPM values were further processed and normalized using downstream analysis tool tximport [92](v1.24.0) in R (v4.2.1)[93].

#### Sample collection, RNA isolation, cDNA synthesis and RT-PCR/qPCR

RNA isolation from drosophila wing discs was performed with the TRIzol® reagent (Invitrogen, Cat. No 15596026), according to manufacturer’s instructions as described previously[21].

Reverse transcription (RT) was carried out with 1μg of total RNA, using random hexamer primers (Invitrogen), OligodT (New England BioLabs, Inc.), RNaseOUT^TM^ Recombinant Ribonuclease Inhibitor (Thermo Fisher Scientific) and Protoscript II reverse transcriptase (New England BioLabs, Inc.), according to manufacturer’s instructions.

Semi-quantitative PCR reactions (RT-PCR) were carried out with 2 µl of cDNA, with the following primers for the Xrp1 DmeEX0007032 splicing event: Xrp1_Exon4_F: GGGAACTTGTGTCGACGACTC and Xrp1_Exon4_R: CCGGACATCGATGACATGTCG. PCR conditions included 30 cycles of denaturation at 94°C for 30’’, annealing at 60°C for 30’’ and extension at 72°C for 90’’. Inclusion/exclusion bands for these events were analyzed by 10% acrylamide electrophoresis and quantified using ImageJ (v1.53t).

Quantitative PCR (qPCR) was performed in a Corbett Rotor Gene™ RG 6000 using Kapa SYBR®FAST (KAPA Biosystems) qPCR mix. The following primer sequences were used: RA_Xrp1_F:CCGGTGTTCATTGATGAATAC, RC_Xrp1_F:CTTTCTAACGGTCAATTAAAAAATACC, Xrp1_R: GTCAAGAATATCCGTCTCAG. As reference gene, aTub84B was used with the following primers: aTub84B_F: CACACCACCCTGGAGCATTC and aTub84B_R: CCAATCAGACGGTTCAGGTTG. qPCR conditions included 30 cycles of denaturation at 94oC for 15’’, annealing at 58oC for 15’’ and extension at 72oC for 15’’.

#### Western blot analysis

Protein lysates from 7 wing imaginal discs of third instar larvae of each genotype were separated on SDS–polyacrylamide gels (12%). After blocking, we incubated with the primary antibody rabbit anti-Xrp1 [26], used the secondary antibody anti-rabbit conjugated with IRDye 680 nm (LI-COR) and imaged on a LI-COR Odyssey scanner system.

#### Proteomic sample preparation using the Sp3-mediated protein digestion protocol

For proteomic analysis we used three to four biological replicas from each genotype. From each genotype wing imaginal discs of third-instar wandering larvae were dissected in 0.1 M sodium phosphate buffer (pH 7.2). We dissected as many discs we could within 10–15 min at room temperature. Drosophila tissue was excised and homogenized in a lysis buffer consisting of 4% SDS, 0.1M DTT, 0.1M TEAB and lysed by incubation at 99 °C for 5 min. Samples were immediately frozen at -80 °C. In total, 50 imaginal discs were used for each biological sample. After thawing, sample were sonicated in waterbath, followed by a centrifugation step for 15 min at 17000xg.

The lysed samples were processed according to the Sp3 protocol (Hughes) including an alkylation step in 200 mM iodoacetamide (Acros Organics). 20 ug of beads (1:1 mixture of hydrophilic and hydrophobic SeraMag carboxylate-modified beads, GE Life Sciences) were added to each sample in 50% ethanol. Protein clean-up was performed on a magnetic rack. The beads were washed two times with 80% ethanol and once with 100% acetonitrile (Fisher Chemical). The captured on beads proteins were digested overnight at 37o C under vigorous shaking (1200 rpm, Eppendorf Thermomixer) with 1 ug Trypsin/LysC (MS grade, Promega) prepared in 25 mM ammonium bicarbonate. Next day, the supernatants were collected and the peptides were purified using a modified Sp3 clean up protocol and finally solubilized in the mobile phase A (0.1% formic acid in water), sonicated and the peptide concentration was determined through absorbance at 280 nm measurement using a nanodrop instrument.

#### LC-MS/MS Analysis

Samples were run on a liquid chromatography tandem mass spectrometry (LC-MS/MS) setup consisting of a Dionex Ultimate 3000RSLC online with a Thermo Q Exactive HF-X Orbitrap mass spectrometer. Peptidic samples were directly injected and separated on an 25 cm-long analytical C18 column (PepSep, 1.9μm3 beads, 75 µm ID) using an 60 minutes long run. A full MS was acquired in profile mode using a Q Exactive HF-X Hybrid Quadrupole-Orbitrap mass spectrometer, operating in the scan range of 375-1400 m/z using 120K resolving power with an AGC of 3x 106 and max IT of 60ms followed by data independent analysis using 8 Th windows (39 loop counts) with 15K resolving power with an AGC of 3x 105 and max IT of 22ms and a normalized collision energy (NCE) of 26. Each biological sample was analyzed in three technical replicas.

#### Proteomic database search

Orbitrap raw data was analyzed in DIA-NN 1.8.1 (Data-Independent Acquisition by Neural Networks)[94] through searching against the reference proteome of *Drosophila melanogaster* (13767 entries) retrieved from Uniprot database in the library free mode of the software, allowing up to two tryptic missed cleavages. A spectral library was created from the DIA runs and used to reanalyse them. DIA-NN default settings have been used with oxidation of methionine residues and acetylation of the protein N-termini set as variable modifications and carbamidomethylation of cysteine residues as fixed modification. N-terminal methionine excision was also enabled. The match between runs (MBR) feature was used for all analyses and the output (precursor) was filtered at 0.01 FDR and finally the protein inference was performed on the level of genes using only proteotypic peptides.

### STATISTICAL ANALYSIS

#### Statistical analyses and plots of qPCR and Alternative Splicing analysis

Statistical tests were performed as indicated in figures legends using R (v4.2.1) or GraphPad Prism (version 8). Principal Component Analysis (PCA) was performed on PSI values of all the exonic splicing events and visualized using ggfortify [95](v0.4.16) and ggplot2 (v3.4.2) packages in R[96] (v4.2.1).

#### Proteomic statistical analysis

For statistical and bioinformatics analysis, as well as for visualization, Perseus, which is part of Maxquant suite, was used[97]. The protein intensities were transformed to logarithmic values [log2(x)]. The protein groups were filtered to obtain at least 2 valid values in at least one group and missing values were imputed from normal distribution. The label-free quantified proteins were subjected to statistical analysis using both ANOVA test and student t-tests, both with permutation-based FDR calculation. The statistical test results were visualized in volcano plot displaying the difference between the two samples expressed as log2(x) versus their statistical significance expressed as −Log10(p-value). Hierarchical clustering was carried out on Z-score transformed LFQ values using average linkage of Euclidian distance. GO Enrichment analysis for biological processes was performed using DAVID functional annotation tools with Flybase Gene ID as identifiers, the Drosophila background and the GOTERM_DIRECT annotation categories [98, 99].

## ACKNOWLEDGMENTS

We specially thank Nicholas Baker and Christos Delidakis for valuable comments on an earlier and the final draft of the manuscript. We thank Chrysoula Pitsouli and Nektarios Tavernarakis for providing feedback on the final draft. We thank Kyriaki Kanakousaki, Maria Lalioti and Konstantina Dimakakou for providing technical support. We thank Christos Delidakis and Erik Storkebaum for flystocks, and Don Rio for the anti-Xrp1 antibody. The research project was supported by the Hellenic Foundation for Research and Innovation (H.F.R.I.) under the “3rd Call for H.F.R.I. Research Projects to support Post-Doctoral Researchers” (Project Number: 7306) to MK.

Confocal microscopy was performed at Fleming using the Leica TCS SP8X WLL confocal system supported by the BIO-IMAGING-GR MIS 5002755 and the Zeiss LSM 900 Airyscan supported by the project MIS 6004752 funded by the Regional Operational Programme ‘ATTICA’ (NSRF 2021-2027) and co-financed by Greece and the European Union (European Regional Development Fund). Proteomic analysis supported by the project pMED-GR (MIS 5002802), which is implemented under the Action “Reinforcement of the Research and Innovation Infrastructure”, funded by the Operational Programme; Competitiveness, Entrepreneurship and Innovation; (NSRF 2014-2020) and co-financed by Greece and the European Union (European Regional Development Fund). Drosophila stocks were obtained from the Bloomington Drosophila Stock Center (supported by NIH P40OD018537), the Vienna Drosophila RNAi Center[100] and FlyORF library[86].

## AUTHOR CONTRIBUTIONS

E.T. and K.K. performed experiments, analyzed and interpreted data. M.P. performed alternative splicing, isoform and RT-PCR/qPCR analysis. M.L. performed experiments. E.M.C. interpreted data. M.S. performed the proteomics experiments and analyzed the data. P.K. supervised alternative splicing analysis, isoform and RT-PCR/qPCR experiments, and analyzed data and edited the manuscript. M.K. conceptualized and led the study, designed and performed experiments, analyzed and interpreted all of the data, acquired funding, and wrote the manuscript.

## INCLUSION AND DIVERSITY

We support inclusive, diverse, and equitable conduct of research.

## SUPPLEMENTAL INFORMATION

Document S1: Figures S1-S7

Supplementary Table 1: PSI values for Xrp1

Supplementary Table 2: Full genotypes for all the experiments, heat shock times and N number for each experiment.

## Supplemental Figures and Legends

**Figure S1.**
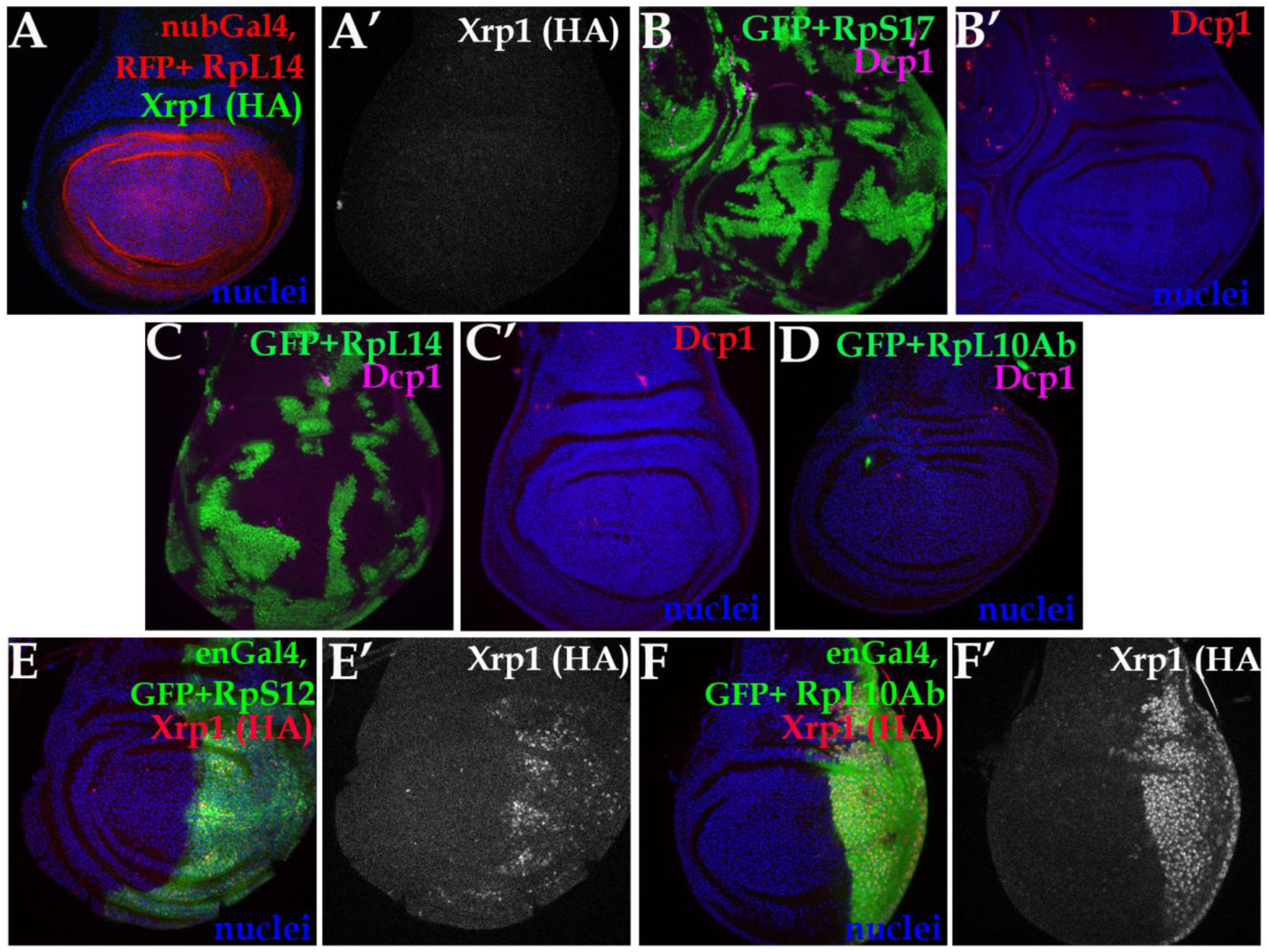
(Related to Figure 1) Confocal images of wing imaginal discs of the indicated genotypes. Panels B-D show projections of Dcp1 staining of the central disc-proper region. All the other panels show single confocal planes. (A) nub-Gal4 over-expression of RFP (red) together with RpL14 in the wing pouch could not induce Xrp1-HA expression. (B,C) Clonal overexpression of GFP (green) together with RpS17 (B) or RpL14 (C) in wild-type discs did not induce boundary cell death (Dcp1, magenta in B,C and red in B’, C’). (D) Clones overexpressing RpL10Ab-GFP (green in D) were never recovered. (E) en-Gal4 drives the co-expression of GFP together with RpS12 in the posterior domain of a wild-type wing disc, which results in the induction of the endogenously HA-tagged Xrp1 allele. (F) en-Gal4 drives the co-expression of GFP together with GFP-RpL10Ab in the posterior domain of a wild-type wing disc, inducing strongly the endogenously HA-tagged Xrp1 allele and reducing the posterior compartment size.

**Figure S2.**
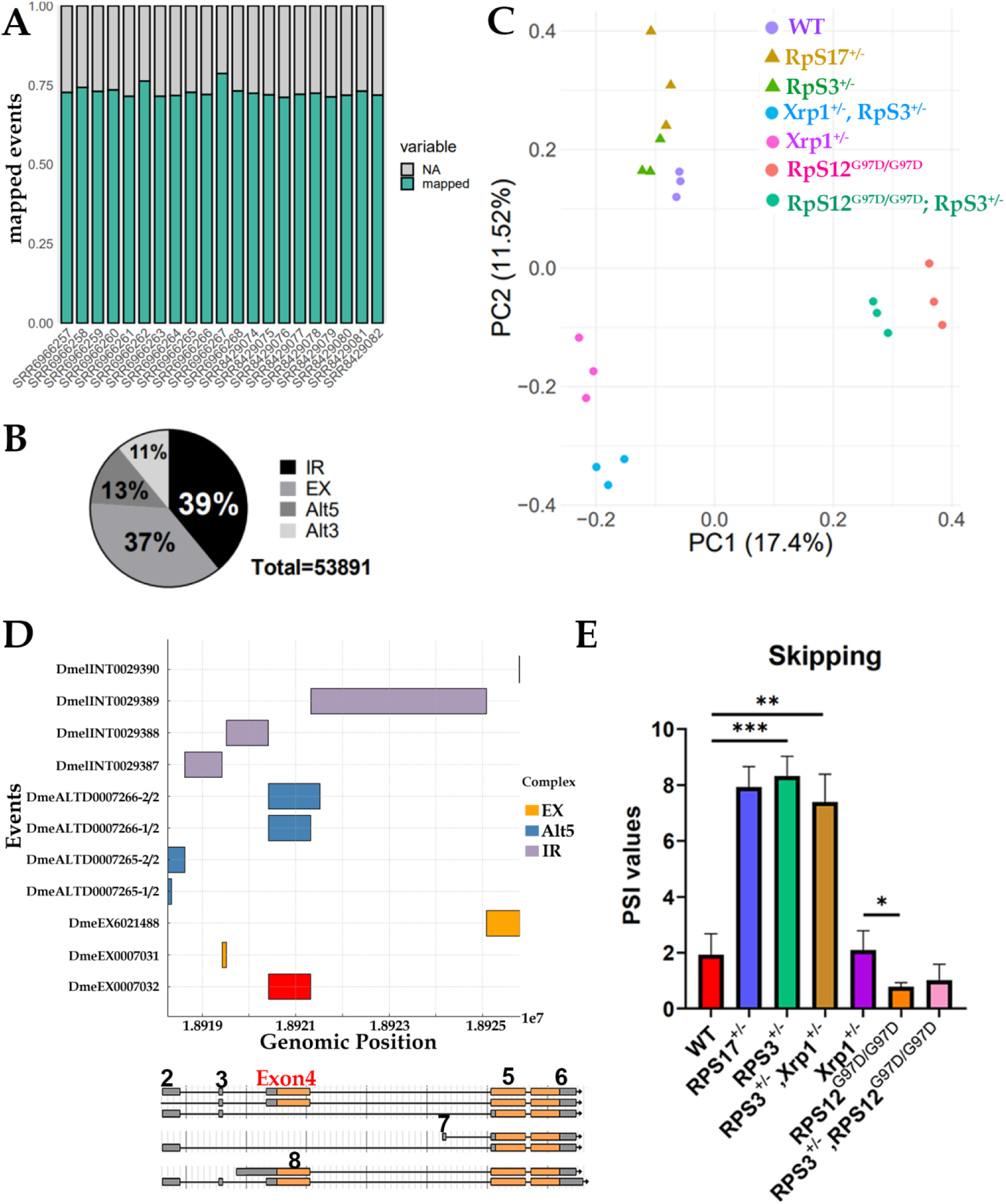
Related to Figure 2. (A) Bar graph showing the percentage of mapped alternative splicing events in all the individual RNA seq samples analyzed. (B) Pie-chart graph of the distribution of the mapped alternative splicing events based on their type. Different shading corresponds to the different event types (IR: Intron Retention; EX: alternative exon; Alt5: alternative 5’ site; Alt3: Alternative 3’ splice site). (C) Principal Component Analysis (PCA) illustrates the distribution of the PSI values for all splicing events mapped across all the genotypes depicted. Each point represents an individual sample analyzed. (D) Bar graph illustrating the genomic locations affected by the annotated alternative splicing events of *Xrp1* (y-axis). The x-axis shows the position on the chr3R chromosome. The y-axis shows the VastdB ID number for each AS event. Events are categorized into three groups based on their type: EX (orange), Alt5 (blue), and intronic or IR (purple). Each event spans specific genomic regions, represented as horizontal bars. The DmeEX0007032 event is highlighted in red for emphasis. *Xrp1* gene locus is presented below for facilitating the identification of the events. (E) PSI values of the *Xrp1* exon 4 skipping from the final mRNA, represented as mean ± SEM. *p-values* are calculated by Student’s t-test (** p< 0.01, *** p< 0.001). (different genotypes presented: wild-type in red, *RpS17*^+/−^ in blue, *RpS3*^+/−^ in green, *Xrp1^+/−^, RpS3^+/−^*in brown, *Xrp1^+/−^* in purple, *RpS12^G97D^*^/G97D^ in orange and *RpS12^G97D^/^G97D^,RpS3^+/−^* in pink

**Figure S3.**
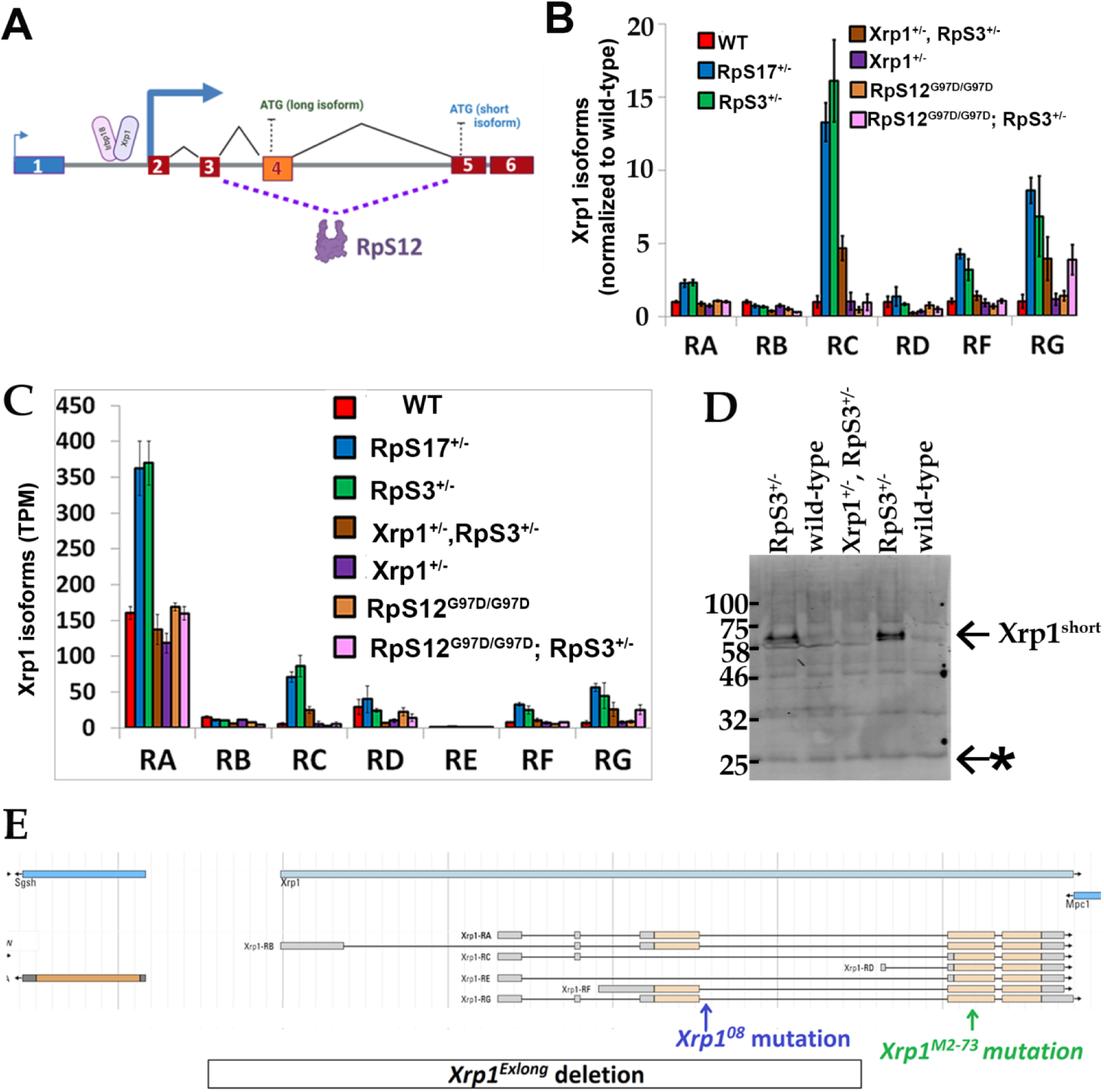
Related to Figure 3. (A) Model showing the RpS12-mediated splicing of Xrp1. Exon 4 skipping results in the production of the RC transcript, encoding the Xrp1^short^ isoform. This isoform, together with is binding partner Irbp18(Blanco et al., 2020), autoregulates Xrp1 in *Minute* cells. Exon 4 contains the Xrp1^long^ translation start site (ATG), whereas the Xrp1^short^ isoform translation initiates within exon 5. (B) Fold change of the Xrp1 mRNA isoforms normalized to wild-type levels. Graph depicts the ratio of transcripts per million of each isoform in indicated genotypes normalized to transcripts per million of each isoform in wild type cells (*wild-type* in red; *RpS17*^+/−^ in blue; *RpS3*^+/−^ in green; *Xrp1^+/−^, RpS3^+/−^* in brown; *Xrp1^+/−^* in purple; *RpS12^G97D/G97D^* in orange; and *RpS12^G97D^/^G97D^, RpS3^+/−^* in pink). Error bars represent ±1 SD derived from RNAseq data. (C) Isoform analysis of Xrp1 transcripts derived from RNAseq data of the genotypes shown in Figure 2. Graph depicts transcripts per million of each of the seven isoforms (RA, RB, RC, RD, RE, RF and RG) in indicated genotypes. (*wild-type* in red; *RpS17*^+/−^ in blue; *RpS3*^+/−^ in green, *Xrp1^+/−^, RpS3^+/−^* in brown; *Xrp1^+/−^*in purple; *RpS12^G97D/G97D^* in orange; and *RpS12^G97D^/^G97D^, RpS3^+/−^* in pink). Error bars represent ±1 SD. (D) Western blots from proteomic extracts of wing discs of the following genotypes: *wild-type*, *RpS3*^+/−^ and *Xrp1^+/−^; RpS3^+/−^*. Only the short isoform is detected by probing with anti-Xrp1 antibody. Of note, the band of the short Xrp1 isoform migrates higher than its expected size [Igaki T, personal communication and (Mallik et al., 2018)]. Xrp1^short^ protein isoform is increased in *RpS3*^+/−^ cells and its induction is abrogated in *Rp*^+/−^ cells heterozygous for the dominant *Xrp1^M2^*^-73^ null allele (*Xrp1*^+/−^). Background bands (asterisk) serve as loading control. (E) Schematic view of chromosome 3R region around the *Xrp1* locus. Locations of the *Xrp1^08^* intronic mutation(Baillon et al., 2018) and *Xrp1^M2^*^-73^ mutation(Lee et al., 2018) are shown. Also the DNA segment that is absent from the deletion strain *Xrp1^Exlong^* (Mallik et al., 2018) is indicated by the box.

**Figure S4.**
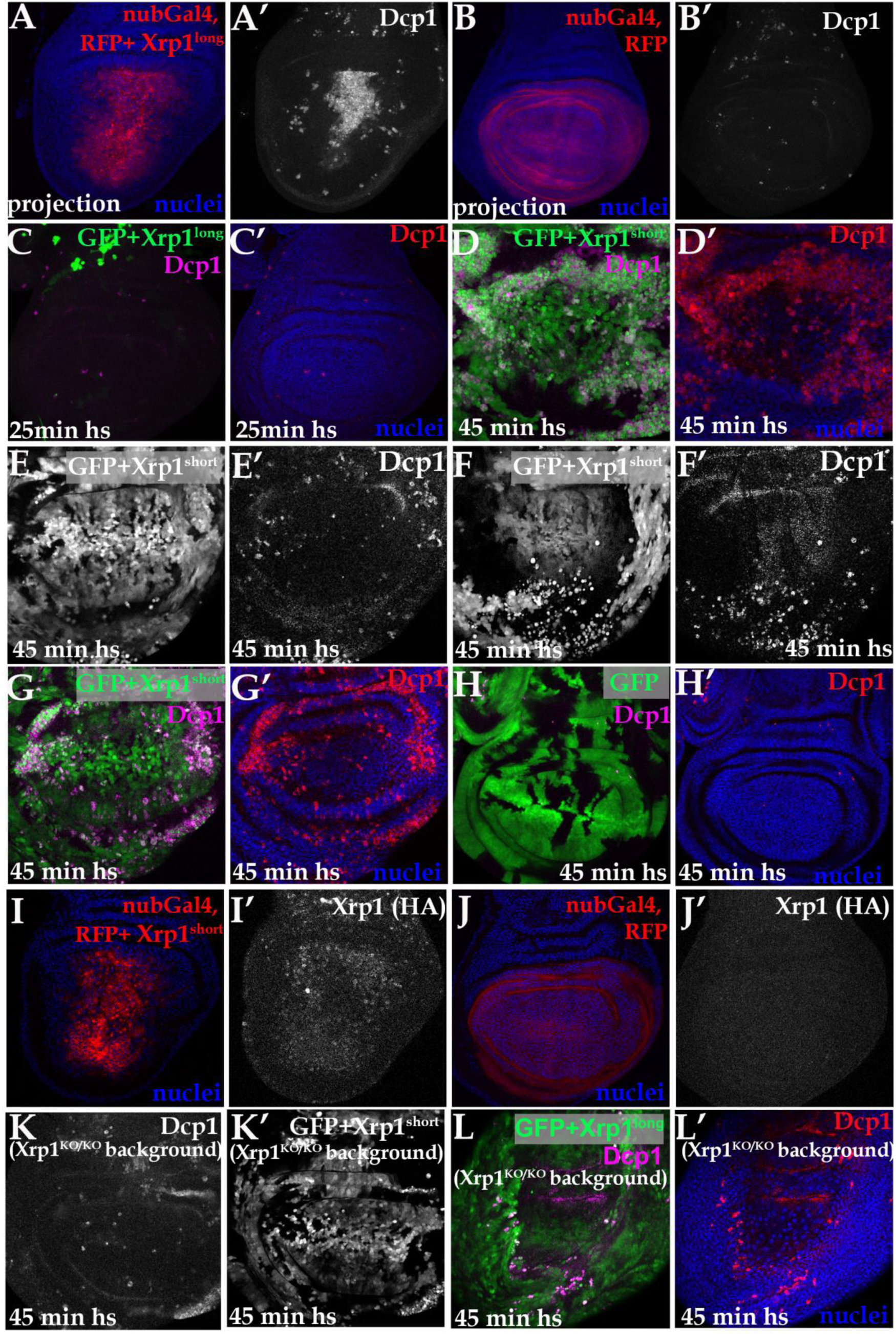
Related to Figure 4. Confocal images of wing imaginal discs of the indicated genotypes. (A-C, K-L) projections of Dcp1 staining of the central disc-proper region. (D-J) single confocal planes. (A) nub-Gal4 drives the co-expression of UAS-RFP reporter (A) together with UAS-Xrp1^long^ in the wing pouch. Xrp1^long^ overexpression resulted in massive cell death (Dcp1, A’) (B) nub-Gal4 drives the expression of UAS-RFP reporter (A) in the wing pouch. Negligible Dcp1 staining is detected (Dcp1, B’). (C) Clonal overexpression of Xrp1^long^ together with a UAS-GFP reporter (green, c) in wing discs did not survive (Dcp1, magenta in C and red in C’). Clones were induced by 25 min heat shock. (D) Cell death at the basal surface of the disc shown in 4C (Dcp1, magenta in D and red in D’). (E, F) Single channels of the disc shown in 4D,E (GFP, E,F) (Dcp1,E’,F’) (G) Cell death at the basal surface of the disc shown in 4D,E (Dcp1, magenta in G and red in G’). (H) Cell death in control clones induced by 45 min heat-shock overexpressing UAS-GFP. (I) nub-Gal4 drives the co-expression of UAS-RFP reporter (I) together with UAS-Xrp1^long^ in the wing pouch, which results in the induction of the endogenously HA-tagged Xrp1 allele (I’) (J) nub-Gal4 drives the expression of UAS-RFP reporter (J) in the wing pouch, which does not causes induction of the endogenously HA-tagged Xrp1 allele (J’) (K) Clonal overexpression of the Xrp1^short^ isoform (GFP channel in K’) in a homozygous null Xrp1^KO^ disc exhibit boundary cell death (Dcp1, K). Clones induced by 45 min heat shock. (L) Clonal overexpression of the Xrp1^long^ isoform (GFP channel in L’) in a heterozygous null Xrp1^KO^ disc exhibit boundary cell death (Dcp1, L). Clones induced by 45 min heat shock.

**Figure S5.**
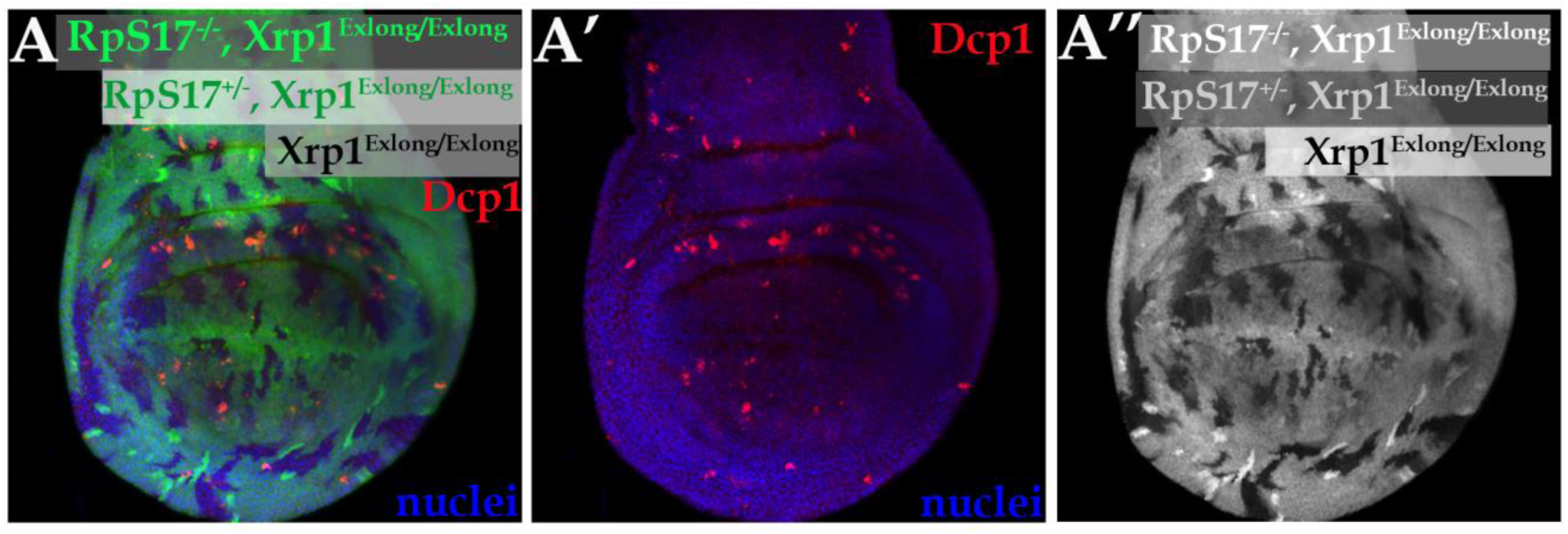
Related to Figure 5. Confocal images of wing imaginal discs of the indicated genotypes. (A) Mosaic wing disc homozygous for the Xrp1^Exlong^ allele, where the RpS17^+/−^; Xrp1^Exlong/Exlong^ cells present boundary cell death at the interfaces with the RpS17^+/+^; Xrp1^Exlong/Exlong^ cells. Small clones of RpS17^-/-^; Xrp1^Exlong/Exlong^ survived 72h after induction (stronger green labeling for ß-galactosidase). In the Xrp1^Exlong^ allele, the two thirds of the Xrp1 gene is deleted, predicted to abolish expression of all of the Xrp1^long^ isoforms. In Xrp1^Exlong^ flies some residual Xrp1 transcript is detected (∼8% of WT levels), possibly reflecting expression of the Xrp1^short^ isoform (Mallik et al., 2018).

**Figure S6.**
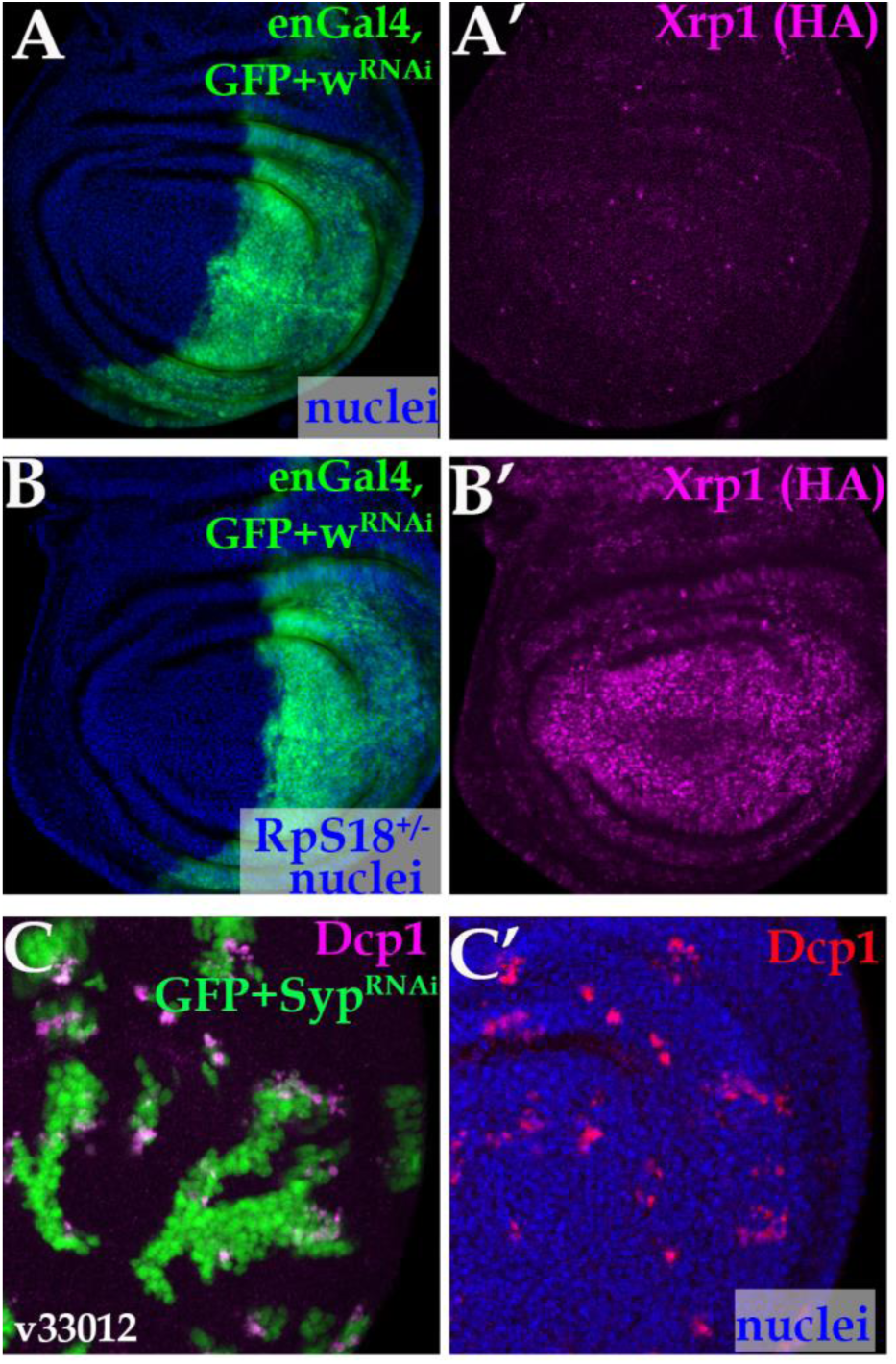
Related to Figure 7. Confocal images of wing imaginal discs of indicated genotypes. A,B, single confocal planes. C, Projection of the central disc-proper region. (A) *white* depletion in the posterior compartment (green in A) of a wild-type wing disc has no effect on Xrp1 expression (magenta in A’). (B) *white* depletion in the posterior compartment (green in B) of an *RpS18*^+/−^ wing disc has no effect on Xrp1 expression (magenta in B’). (C) Clonal *Syp* depletion by an RNAi line (v33012) (green in C) in a wild-type disc induces cell death (active Dcp1 staining, magenta in C and red in C’) at clone boundaries.

**Figure S7.**
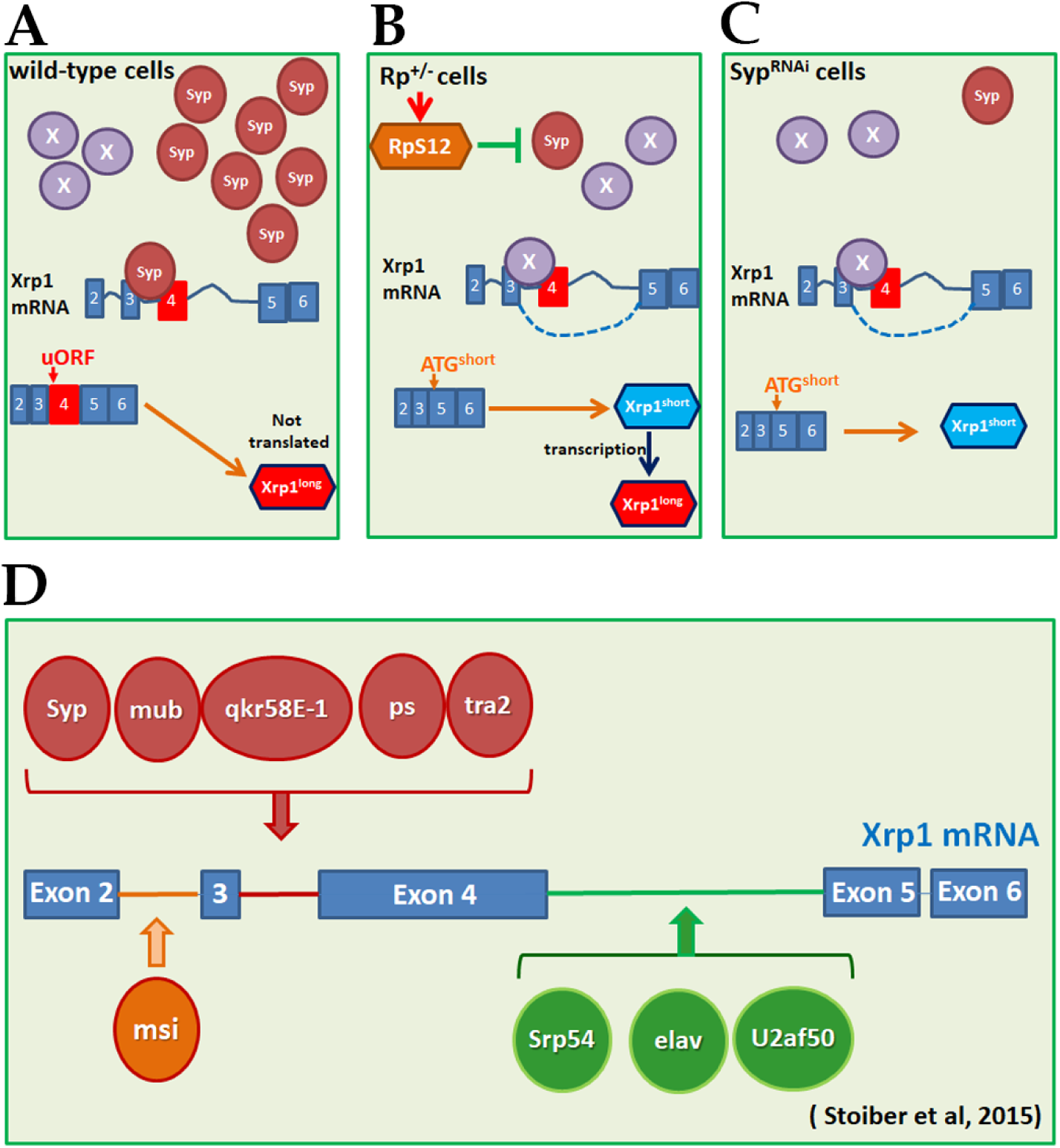
Schematic models. (A) In wild-type cells, Syncrip negatively regulates Xrp1 expression, by preventing the binding of another RBP that promotes *Xrp1* exon 4 skipping. Therefore, Syncrip binding results in the *Xrp1^long^* RA transcript being the predominant isoform. The presence of uORFs in the 5’ UTR of this *Xrp1^long^* RA transcript limits its efficient translation. (B) In *Rp*^+/−^ cells, RpS12 reduces Syncrip levels. This allows a competitive RBP, X, to promote Xrp1 exon 4 skipping, producing the Xrp1^short^ RC transcript, which is efficiently translated. Xrp1^short^ protein isoform, by autoregulation, increases the levels of both isoforms. (C) Syp depletion in wild-type cells mimics the *Minute* condition, allowing a competing RBP, X, to promote exon 4 skipping and produce the *Xrp1^short^* RC transcript, which is efficiently translated. (D) Schematic of the Xrp1 transcript showing the highest affinity binding sites of RNA-binding proteins identified by Stoiber et al. (2015)(Stoiber et al., 2015).

## REFERENCES

1. Narla, A., and Ebert, B.L. (2010). Ribosomopathies: human disorders of ribosome dysfunction. Blood 115, 3196–3205.

2. Ulirsch, J.C., Verboon, J.M., Kazerounian, S., Guo, M.H., Yuan, D., Ludwig, L.S., Handsaker, R.E., Abdulhay, N.J., Fiorini, C., Genovese, G., et al. (2018). The Genetic Landscape of Diamond-Blackfan Anemia. Am J Hum Genet 103, 930–947.

3. Aspesi, A., and Ellis, S.R. (2019). Rare ribosomopathies: insights into mechanisms of cancer. Nat Rev Cancer.

4. Sulima, S.O., Hofman, I.J.F., De Keersmaecker, K., and Dinman, J.D. (2017). How Ribosomes Translate Cancer. Cancer Discov 7, 1069–1087.

5. Marygold, S.J., Roote, J., Reuter, G., Lambertsson, A., Ashburner, M., Millburn, G.H., Harrison, P.M., Yu, Z., Kenmochi, N., Kaufman, T.C., et al. (2007). The ribosomal protein genes and Minute loci of Drosophila melanogaster. Genome Biol 8, R216.

6. Morata, G., and Ripoll, P. (1975). Minutes: mutants of drosophila autonomously affecting cell division rate. Dev Biol 42, 211–221.

7. Li, W., and Baker, N.E. (2007). Engulfment is required for cell competition. Cell 129, 1215–1225.

8. Moreno, E., Basler, K., and Morata, G. (2002). Cells compete for decapentaplegic survival factor to prevent apoptosis in Drosophila wing development. Nature 416, 755–759.

9. Kiparaki, M., and Baker, N.E. (2023). Ribosomal protein mutations and cell competition: autonomous and nonautonomous effects on a stress response. Genetics 224.

10. Morata, G. (2021). Cell competition: A historical perspective. Dev Biol.

11. Simpson, P., and Morata, G. (1981). Differential mitotic rates and patterns of growth in compartments in the Drosophila wing. Dev Biol 85, 299–308.

12. Baker, N.E. (2020). Emerging mechanisms of cell competition. Nat Rev Genet.

13. Bondar, T., and Medzhitov, R. (2010). p53-mediated hematopoietic stem and progenitor cell competition. Cell Stem Cell 6, 309–322.

14. Ellis, S.J., Gomez, N.C., Levorse, J., Mertz, A.F., Ge, Y., and Fuchs, E. (2019). Distinct modes of cell competition shape mammalian tissue morphogenesis. Nature 569, 497–502.

15. Levayer, R., and Moreno, E. (2013). Mechanisms of cell competition: themes and variations. J Cell Biol 200, 689–698.

16. Bowling, S., Lawlor, K., and Rodriguez, T.A. (2019). Cell competition: the winners and losers of fitness selection. Development 146.

17. Claveria, C., and Torres, M. (2016). Cell Competition: Mechanisms and Physiological Roles. Annu Rev Cell Dev Biol 32, 411–439.

18. Vishwakarma, M., and Piddini, E. (2020). Outcompeting cancer. Nat Rev Cancer 20, 187–198.

19. van Neerven, S.M., and Vermeulen, L. (2023). Cell competition in development, homeostasis and cancer. Nat Rev Mol Cell Biol 24, 221–236.

20. Colom, B., Herms, A., Hall, M.W.J., Dentro, S.C., King, C., Sood, R.K., Alcolea, M.P., Piedrafita, G., Fernandez-Antoran, D., Ong, S.H., et al. (2021). Mutant clones in normal epithelium outcompete and eliminate emerging tumours. Nature 598, 510–514.

21. Lee, C.H., Kiparaki, M., Blanco, J., Folgado, V., Ji, Z., Kumar, A., Rimesso, G., and Baker, N.E. (2018). A Regulatory Response to Ribosomal Protein Mutations Controls Translation, Growth, and Cell Competition. Dev Cell 46, 456–469 e454.

22. Baillon, L., Germani, F., Rockel, C., Hilchenbach, J., and Basler, K. (2018). Xrp1 is a transcription factor required for cell competition-driven elimination of loser cells. Sci Rep 8, 17712.

23. Lee, C.H., Rimesso, G., Reynolds, D.M., Cai, J., and Baker, N.E. (2016). Whole-Genome Sequencing and iPLEX MassARRAY Genotyping Map an EMS-Induced Mutation Affecting Cell Competition in Drosophila melanogaster. G3 (Bethesda) 6, 3207–3217.

24. Blanco, J., Cooper, J.C., and Baker, N.E. (2020). Roles of C/EBP class bZip proteins in the growth and cell competition of Rp (’Minute’) mutants in Drosophila. Elife 9.

25. Kiparaki, M., Khan, C., Folgado-Marco, V., Chuen, J., Moulos, P., and Baker, N.E. (2022). The transcription factor Xrp1 orchestrates both reduced translation and cell competition upon defective ribosome assembly or function. Elife 11.

26. Francis, M.J., Roche, S., Cho, M.J., Beall, E., Min, B., Panganiban, R.P., and Rio, D.C. (2016). Drosophila IRBP bZIP heterodimer binds P-element DNA and affects hybrid dysgenesis. Proc Natl Acad Sci U S A 113, 13003–13008.

27. Ji, Z., Kiparaki, M., Folgado, V., Kumar, A., Blanco, J., Rimesso, G., Chuen, J., Liu, Y., Zheng, D., and Baker, N.E. (2019). Drosophila RpS12 controls translation, growth, and cell competition through Xrp1. PLoS Genet 15, e1008513.

28. Kale, A., Ji, Z., Kiparaki, M., Blanco, J., Rimesso, G., Flibotte, S., and Baker, N.E. (2018). Ribosomal Protein S12e Has a Distinct Function in Cell Competition. Dev Cell 44, 42–55 e44.

29. Martin-Villanueva, S., Galmozzi, C.V., Ruger-Herreros, C., Kressler, D., and de la Cruz, J. (2024). The Beak of Eukaryotic Ribosomes: Life, Work and Miracles. Biomolecules 14.

30. Tyler, D.M., Li, W., Zhuo, N., Pellock, B., and Baker, N.E. (2007). Genes affecting cell competition in Drosophila. Genetics 175, 643–657.

31. Langton, P.F., Baumgartner, M.E., Logeay, R., and Piddini, E. (2021). Xrp1 and Irbp18 trigger a feed-forward loop of proteotoxic stress to induce the loser status. PLoS Genet 17, e1009946.

32. Ochi, N., Nakamura, M., Nagata, R., Wakasa, N., Nakano, R., and Igaki, T. (2021). Cell competition is driven by Xrp1-mediated phosphorylation of eukaryotic initiation factor 2alpha. PLoS Genet 17, e1009958.

33. Kumar, A., and Baker, N.E. (2022). The CRL4 E3 ligase Mahjong/DCAF1 controls cell competition through the transcription factor Xrp1, independently of polarity genes. Development 149.

34. Stankovic, D., Tain, L.S., and Uhlirova, M. (2024). Xrp1 governs the stress response program to spliceosome dysfunction. Nucleic Acids Res 52, 2093–2111.

35. Xue, M., Cong, F., Zheng, W., Xu, R., Liu, X., Bao, H., Sung, Y.Y., Xi, Y., He, F., Ma, J., et al. (2023). Loss of Paip1 causes translation reduction and induces apoptotic cell death through ISR activation and Xrp1. Cell Death Discov 9, 288.

36. Brown, B., Mitra, S., Roach, F.D., Vasudevan, D., and Ryoo, H.D. (2021). The transcription factor Xrp1 is required for PERK-mediated antioxidant gene induction in Drosophila. Elife 10.

37. Destefanis, F., Manara, V., Santarelli, S., Zola, S., Brambilla, M., Viola, G., Maragno, P., Signoria, I., Viero, G., Pasini, M.E., et al. (2022). Reduction of nucleolar NOC1 leads to the accumulation of pre-rRNAs and induces Xrp1, affecting growth and resulting in cell competition. J Cell Sci 135.

38. Boulan, L., Andersen, D., Colombani, J., Boone, E., and Leopold, P. (2019). Inter-Organ Growth Coordination Is Mediated by the Xrp1-Dilp8 Axis in Drosophila. Dev Cell 49, 811–818 e814.

39. Ji, Z., Chuen, J., Kiparaki, M., and Baker, N. (2021). Cell competition removes segmental aneuploid cells from Drosophila imaginal disc-derived tissues based on ribosomal protein gene dose. Elife 10.

40. Baumgartner, M.E., Dinan, M.P., Langton, P.F., Kucinski, I., and Piddini, E. (2021). Proteotoxic stress is a driver of the loser status and cell competition. Nat Cell Biol 23, 136–146.

41. Recasens-Alvarez, C., Alexandre, C., Kirkpatrick, J., Nojima, H., Huels, D.J., Snijders, A.P., and Vincent, J.P. (2021). Ribosomopathy-associated mutations cause proteotoxic stress that is alleviated by TOR inhibition. Nat Cell Biol 23, 127–135.

42. Albert, B., Kos-Braun, I.C., Henras, A.K., Dez, C., Rueda, M.P., Zhang, X., Gadal, O., Kos, M., and Shore, D. (2019). A ribosome assembly stress response regulates transcription to maintain proteome homeostasis. Elife 8.

43. Tye, B.W., Commins, N., Ryazanova, L.V., Wuhr, M., Springer, M., Pincus, D., and Churchman, L.S. (2019). Proteotoxicity from aberrant ribosome biogenesis compromises cell fitness. Elife 8.

44. Thomas, A., Lee, P.J., Dalton, J.E., Nomie, K.J., Stoica, L., Costa-Mattioli, M., Chang, P., Nuzhdin, S., Arbeitman, M.N., and Dierick, H.A. (2012). A versatile method for cell-specific profiling of translated mRNAs in Drosophila. PLoS One 7, e40276.

45. Gramates, L.S., Agapite, J., Attrill, H., Calvi, B.R., Crosby, M.A., Dos Santos, G., Goodman, J.L., Goutte-Gattat, D., Jenkins, V.K., Kaufman, T., et al. (2022). FlyBase: a guided tour of highlighted features. Genetics 220.

46. Stoiber, M.H., Olson, S., May, G.E., Duff, M.O., Manent, J., Obar, R., Guruharsha, K.G., Bickel, P.J., Artavanis-Tsakonas, S., Brown, J.B., et al. (2015). Extensive cross-regulation of post-transcriptional regulatory networks in Drosophila. Genome Res 25, 1692–1702.

47. Singh, A.K., Choudhury, S.R., De, S., Zhang, J., Kissane, S., Dwivedi, V., Ramanathan, P., Petric, M., Orsini, L., Hebenstreit, D., et al. (2019). The RNA helicase UPF1 associates with mRNAs co-transcriptionally and is required for the release of mRNAs from gene loci. Elife 8.

48. Young, S.K., and Wek, R.C. (2016). Upstream Open Reading Frames Differentially Regulate Gene-specific Translation in the Integrated Stress Response. J Biol Chem 291, 16927–16935.

49. Tsurui-Nishimura, N., Nguyen, T.Q., Katsuyama, T., Minami, T., Furuhashi, H., Oshima, Y., and Kurata, S. (2013). Ectopic antenna induction by overexpression of CG17836/Xrp1 encoding an AT-hook DNA binding motif protein in Drosophila. Biosci Biotechnol Biochem 77, 339–344.

50. Menendez, J., Perez-Garijo, A., Calleja, M., and Morata, G. (2010). A tumor-suppressing mechanism in Drosophila involving cell competition and the Hippo pathway. Proc Natl Acad Sci U S A 107, 14651–14656.

51. Kucinski, I., Dinan, M., Kolahgar, G., and Piddini, E. (2017). Chronic activation of JNK JAK/STAT and oxidative stress signalling causes the loser cell status. Nat Commun 8, 136.

52. Mallik, M., Catinozzi, M., Hug, C.B., Zhang, L., Wagner, M., Bussmann, J., Bittern, J., Mersmann, S., Klambt, C., Drexler, H.C.A., et al. (2018). Xrp1 genetically interacts with the ALS-associated FUS orthologue caz and mediates its toxicity. J Cell Biol 217, 3947–3964.

53. Kale, A., Li, W., Lee, C.H., and Baker, N.E. (2015). Apoptotic mechanisms during competition of ribosomal protein mutant cells: roles of the initiator caspases Dronc and Dream/Strica. Cell Death Differ 22, 1300–1312.

54. Dassi, E. (2017). Handshakes and Fights: The Regulatory Interplay of RNA-Binding Proteins. Front Mol Biosci 4, 67.

55. Kampen, K.R., Sulima, S.O., Vereecke, S., and De Keersmaecker, K. (2020). Hallmarks of ribosomopathies. Nucleic Acids Res 48, 1013–1028.

56. De Keersmaecker, K., Atak, Z.K., Li, N., Vicente, C., Patchett, S., Girardi, T., Gianfelici, V., Geerdens, E., Clappier, E., Porcu, M., et al. (2013). Exome sequencing identifies mutation in CNOT3 and ribosomal genes RPL5 and RPL10 in T-cell acute lymphoblastic leukemia. Nat Genet 45, 186–190.

57. Vlachos, A., Rosenberg, P.S., Atsidaftos, E., Alter, B.P., and Lipton, J.M. (2012). Incidence of neoplasia in Diamond Blackfan anemia: a report from the Diamond Blackfan Anemia Registry. Blood 119, 3815–3819.

58. Ajore, R., Raiser, D., McConkey, M., Joud, M., Boidol, B., Mar, B., Saksena, G., Weinstock, D.M., Armstrong, S., Ellis, S.R., et al. (2017). Deletion of ribosomal protein genes is a common vulnerability in human cancer, especially in concert with TP53 mutations. EMBO Mol Med 9, 498–507.

59. Mills, E.W., and Green, R. (2017). Ribosomopathies: There’s strength in numbers. Science 358.

60. Yang, Z., Keel, S.B., Shimamura, A., Liu, L., Gerds, A.T., Li, H.Y., Wood, B.L., Scott, B.L., and Abkowitz, J.L. (2016). Delayed globin synthesis leads to excess heme and the macrocytic anemia of Diamond Blackfan anemia and del(5q) myelodysplastic syndrome. Sci Transl Med 8, 338ra367.

61. Dutt, S., Narla, A., Lin, K., Mullally, A., Abayasekara, N., Megerdichian, C., Wilson, F.H., Currie, T., Khanna-Gupta, A., Berliner, N., et al. (2011). Haploinsufficiency for ribosomal protein genes causes selective activation of p53 in human erythroid progenitor cells. Blood 117, 2567–2576.

62. Brodsky, M.H., Weinert, B.T., Tsang, G., Rong, Y.S., McGinnis, N.M., Golic, K.G., Rio, D.C., and Rubin, G.M. (2004). Drosophila melanogaster MNK/Chk2 and p53 regulate multiple DNA repair and apoptotic pathways following DNA damage. Mol Cell Biol 24, 1219–1231.

63. Akdemir, F., Christich, A., Sogame, N., Chapo, J., and Abrams, J.M. (2007). p53 directs focused genomic responses in Drosophila. Oncogene 26, 5184–5193.

64. Baker, N.E., Kiparaki, M., and Khan, C. (2019). A potential link between p53, cell competition and ribosomopathy in mammals and in Drosophila. Dev Biol 446, 17–19.

65. Zhou, X., Liao, W.J., Liao, J.M., Liao, P., and Lu, H. (2015). Ribosomal proteins: functions beyond the ribosome. J Mol Cell Biol 7, 92–104.

66. Chen, J., Crutchley, J., Zhang, D., Owzar, K., and Kastan, M.B. (2017). Identification of a DNA Damage-Induced Alternative Splicing Pathway That Regulates p53 and Cellular Senescence Markers. Cancer Discov 7, 766–781.

67. Yanagi, S., Iida, T., and Kobayashi, T. (2022). RPS12 and UBC4 Are Related to Senescence Signal Production in the Ribosomal RNA Gene Cluster. Mol Cell Biol 42, e0002822.

68. Warner, J.R., and McIntosh, K.B. (2009). How common are extraribosomal functions of ribosomal proteins? Mol Cell 34, 3–11.

69. Rajan, K.S., Madmoni, H., Bashan, A., Taoka, M., Aryal, S., Nobe, Y., Doniger, T., Galili Kostin, B., Blumberg, A., Cohen-Chalamish, S., et al. (2023). A single pseudouridine on rRNA regulates ribosome structure and function in the mammalian parasite Trypanosoma brucei. Nat Commun 14, 7462.

70. Genuth, N.R., and Barna, M. (2018). Heterogeneity and specialized functions of translation machinery: from genes to organisms. Nat Rev Genet 19, 431–452.

71. Hashem, Y., des Georges, A., Dhote, V., Langlois, R., Liao, H.Y., Grassucci, R.A., Hellen, C.U., Pestova, T.V., and Frank, J. (2013). Structure of the mammalian ribosomal 43S preinitiation complex bound to the scanning factor DHX29. Cell 153, 1108–1119.

72. Brumwell, A., Fell, L., Obress, L., and Uniacke, J. (2020). Hypoxia influences polysome distribution of human ribosomal protein S12 and alternative splicing of ribosomal protein mRNAs. RNA 26, 361–371.

73. Martin-Villanueva, S., Fernandez-Fernandez, J., Rodriguez-Galan, O., Fernandez-Boraita, J., Villalobo, E., and de La Cruz, J. (2020). Role of the 40S beak ribosomal protein eS12 in ribosome biogenesis and function in Saccharomyces cerevisiae. RNA Biol 17, 1261–1276.

74. Kurylo, C.M., Parks, M.M., Juette, M.F., Zinshteyn, B., Altman, R.B., Thibado, J.K., Vincent, C.T., and Blanchard, S.C. (2018). Endogenous rRNA Sequence Variation Can Regulate Stress Response Gene Expression and Phenotype. Cell Rep 25, 236–248 e236.

75. Takei, S., Togo-Ohno, M., Suzuki, Y., and Kuroyanagi, H. (2016). Evolutionarily conserved autoregulation of alternative pre-mRNA splicing by ribosomal protein L10a. Nucleic Acids Res 44, 5585–5596.

76. Shi, Z., Fujii, K., Kovary, K.M., Genuth, N.R., Rost, H.L., Teruel, M.N., and Barna, M. (2017). Heterogeneous Ribosomes Preferentially Translate Distinct Subpools of mRNAs Genome-wide. Mol Cell 67, 71–83 e77.

77. Jia, L., Mao, Y., Ji, Q., Dersh, D., Yewdell, J.W., and Qian, S.B. (2020). Decoding mRNA translatability and stability from the 5’ UTR. Nat Struct Mol Biol 27, 814–821.

78. Boutros, M., Kiger, A.A., Armknecht, S., Kerr, K., Hild, M., Koch, B., Haas, S.A., Paro, R., and Perrimon, N. (2004). Genome-wide RNAi analysis of growth and viability in Drosophila cells. Science 303, 832–835.

79. Mizutani, A., Fukuda, M., Ibata, K., Shiraishi, Y., and Mikoshiba, K. (2000). SYNCRIP, a cytoplasmic counterpart of heterogeneous nuclear ribonucleoprotein R, interacts with ubiquitous synaptotagmin isoforms. J Biol Chem 275, 9823–9831.

80. Titlow, J., Robertson, F., Jarvelin, A., Ish-Horowicz, D., Smith, C., Gratton, E., and Davis, I. (2020). Syncrip/hnRNP Q is required for activity-induced Msp300/Nesprin-1 expression and new synapse formation. J Cell Biol 219.

81. Lee, J.Y., Huang, N., Samuels, T.J., and Davis, I. (2025). Imp/IGF2BP and Syp/SYNCRIP temporal RNA interactomes uncover combinatorial networks of regulators of Drosophila brain development. Sci Adv 11, eadr6682.

82. Bannai, H., Fukatsu, K., Mizutani, A., Natsume, T., Iemura, S., Ikegami, T., Inoue, T., and Mikoshiba, K. (2004). An RNA-interacting protein, SYNCRIP (heterogeneous nuclear ribonuclear protein Q1/NSAP1) is a component of mRNA granule transported with inositol 1,4,5-trisphosphate receptor type 1 mRNA in neuronal dendrites. J Biol Chem 279, 53427–53434.

83. Neubauer, G., King, A., Rappsilber, J., Calvio, C., Watson, M., Ajuh, P., Sleeman, J., Lamond, A., and Mann, M. (1998). Mass spectrometry and EST-database searching allows characterization of the multi-protein spliceosome complex. Nat Genet 20, 46–50.

84. Herrejon Chavez, F., Luo, H., Cifani, P., Pine, A., Chu, E.L., Joshi, S., Barin, E., Schurer, A., Chan, M., Chang, K., et al. (2023). RNA binding protein SYNCRIP maintains proteostasis and self-renewal of hematopoietic stem and progenitor cells. Nat Commun 14, 2290.

85. Catinozzi, M., Mallik, M., Frickenhaus, M., Been, M., Sijlmans, C., Kulshrestha, D., Alexopoulos, I., Weitkunat, M., Schnorrer, F., and Storkebaum, E. (2020). The Drosophila FUS ortholog cabeza promotes adult founder myoblast selection by Xrp1-dependent regulation of FGF signaling. PLoS Genet 16, e1008731.

86. Bischof, J., Bjorklund, M., Furger, E., Schertel, C., Taipale, J., and Basler, K. (2013). A versatile platform for creating a comprehensive UAS-ORFeome library in Drosophila. Development 140, 2434–2442.

87. Golic, K.G., and Lindquist, S. (1989). The FLP recombinase of yeast catalyzes site-specific recombination in the Drosophila genome. Cell 59, 499–509.

88. Xu, T., and Rubin, G.M. (1993). Analysis of genetic mosaics in developing and adult Drosophila tissues. Development 117, 1223–1237.

89. Tapial, J., Ha, K.C.H., Sterne-Weiler, T., Gohr, A., Braunschweig, U., Hermoso-Pulido, A., Quesnel-Vallieres, M., Permanyer, J., Sodaei, R., Marquez, Y., et al. (2017). An atlas of alternative splicing profiles and functional associations reveals new regulatory programs and genes that simultaneously express multiple major isoforms. Genome Res 27, 1759–1768.

90. Irimia, M., Weatheritt, R.J., Ellis, J.D., Parikshak, N.N., Gonatopoulos-Pournatzis, T., Babor, M., Quesnel-Vallieres, M., Tapial, J., Raj, B., O’Hanlon, D., et al. (2014). A highly conserved program of neuronal microexons is misregulated in autistic brains. Cell 159, 1511–1523.

91. Patro, R., Duggal, G., Love, M.I., Irizarry, R.A., and Kingsford, C. (2017). Salmon provides fast and bias-aware quantification of transcript expression. Nat Methods 14, 417–419.

92. Soneson, C., Love, M.I., and Robinson, M.D. (2015). Differential analyses for RNA-seq: transcript-level estimates improve gene-level inferences. F1000Res 4, 1521.

93. R Core Team (2021). R: A Language and Environment for Statistical Computing. Vienna, Austria: R foundation for Statistical Computing. url: https://www.R-project.org/.

94. Demichev, V., Messner, C.B., Vernardis, S.I., Lilley, K.S., and Ralser, M. (2020). DIA-NN: neural networks and interference correction enable deep proteome coverage in high throughput. Nat Methods 17, 41–44.

95. Tang, Y., Horikoshi, M., and Li, W. (2016). ggfortify: Unified interface to visualize statistical results of popular R packages. The R Journal 8, 474–485.

96. Wickham, H. (2016). ggplot2: Elegant Graphics for Data Analysis. Springer-VerlagNew York.

97. Tyanova, S., Temu, T., Sinitcyn, P., Carlson, A., Hein, M.Y., Geiger, T., Mann, M., and Cox, J. (2016). The Perseus computational platform for comprehensive analysis of (prote)omics data. Nat Methods 13, 731–740.

98. Huang da, W., Sherman, B.T., and Lempicki, R.A. (2009). Systematic and integrative analysis of large gene lists using DAVID bioinformatics resources. Nat Protoc 4, 44–57.

99. Sherman, B.T., Hao, M., Qiu, J., Jiao, X., Baseler, M.W., Lane, H.C., Imamichi, T., and Chang, W. (2022). DAVID: a web server for functional enrichment analysis and functional annotation of gene lists (2021 update). Nucleic Acids Res 50, W216–W221.

100. Dietzl, G., Chen, D., Schnorrer, F., Su, K.C., Barinova, Y., Fellner, M., Gasser, B., Kinsey, K., Oppel, S., Scheiblauer, S., et al. (2007). A genome-wide transgenic RNAi library for conditional gene inactivation in Drosophila. Nature 448, 151–156.

