## Supplementary Table 1 for "RpS12-mediated induction of the Xrp1^short^ isoform links ribosomal protein mutations to cell competition"

| GENE | EVENT | COORD | LENGTH | FullCO | COMPLEX |
|---|---|---|---|---|---|
| *strain* | | | | | |
| Xrp1 | DmeEX0007032 | chr3R:18920413-18921319 | 907 | chr3R:18919512,18920413-18921319,18925078 | EX |
| Xrp1 | DmeEX0007031 | chr3R:18919423-18919512 | 90 | chr3R:18918625,18919423-18919512,18920413 | EX |
| Xrp1 | DmeALTD0007265-1/2 | chr3R:18918260-18918341 | 0 | chr3R:18918260-18918341+18918625,18919423 | Alt5 |
| Xrp1 | DmeALTD0007265-2/2 | chr3R:18918260-18918625 | 284 | chr3R:18918260-18918341+18918625,18919423 | Alt5 |
| Xrp1 | DmeALTD0007266-1/2 | chr3R:18920413-18921319 | 0 | chr3R:18920413-18921319+18921518,18925078 | Alt5 |
| Xrp1 | DmeALTD0007266-2/2 | chr3R:18920413-18921518 | 199 | chr3R:18920413-18921319+18921518,18925078 | Alt5 |
| Xrp1 | DmeEX6021488 | chr3R:18925078-18925796 | 719 | chr3R:18921319,18925078-18925796,18925905 | EX |
| Xrp1 | DmeINT0029387 | chr3R:18918626-18919422 | 797 | chr3R:18918260-18918625=18919423-18919512:+ | IR |
| Xrp1 | DmeINT0029388 | chr3R:18919513-18920412 | 900 | chr3R:18919423-18919512=18920413-18921319:+ | IR |
| Xrp1 | DmeINT0029389 | chr3R:18921320-18925077 | 3758 | chr3R:18920413-18921319=18925078-18925796:+ | IR |
| Xrp1 | DmeINT0029390 | chr3R:18925797-18925904 | 108 | chr3R:18925078-18925796=18925905-18926849:+ | IR |

| SCAFFOLD | START | END | STRAND | UPSTRM_EX_BORDER | DOSTRM_EX_BORDER | PSI_SRR6966257 | PSI_SRR6966258 | PSI_SRR6966259 |
|---|---|---|---|---|---|---|---|---|
| | | | | | | *wild-type* | | |
| chr3R | 18920413 | 18921319 | + | 18919512 | 18925078 | 98.35 | 97.23 | 98.67 |
| chr3R | 18919423 | 18919512 | + | 18918625 | 18920413 | 100.00 | 99.53 | 100.00 |
| chr3R | 18918341 | 18918341 | + | 18918260 | 18919423 | 0.21 | 0.00 | 0.53 |
| chr3R | 18918341 | 18918625 | + | 18918260 | 18919423 | 99.79 | 100.00 | 99.47 |
| chr3R | 18921319 | 18921319 | + | 18920413 | 18925078 | 100.00 | 100.00 | 100.00 |
| chr3R | 18921319 | 18921518 | + | 18920413 | 18925078 | 0.00 | 0.00 | 0.00 |
| chr3R | 18925078 | 18925796 | + | 18921319 | 18925905 | 100.00 | 100.00 | 100.00 |
| chr3R | 18918626 | 18919422 | + | 18918260 | 18919512 | 0.00 | 0.00 | 0.13 |
| chr3R | 18919513 | 18920412 | + | 18919423 | 18921319 | 3.09 | 3.05 | 4.19 |
| chr3R | 18921320 | 18925077 | + | 18920413 | 18925796 | 1.33 | 2.01 | 2.05 |
| chr3R | 18925797 | 18925904 | + | 18925078 | 18926849 | 2.44 | 2.29 | 3.76 |

| PSI_SRR6966260 | PSI_SRR6966261 | PSI_SRR6966262 | PSI_SRR6966263 | PSI_SRR6966264 | PSI_SRR6966265 | PSI_SRR6966266 | PSI_SRR6966267 |
|---|---|---|---|---|---|---|---|
| *RpS17* $^{+/-}$ | | | *RpS3* $^{+/-}$ | | | *Xrp1* $^{+/-}$ , *RpS3* $^{+/}$ | |
| 91.24 | 92.39 | 92.60 | 90.87 | 92.01 | 92.16 | 91.62 | 92.60 |
| 99.59 | 99.81 | 99.69 | 99.52 | 99.76 | 99.63 | 100.00 | 99.50 |
| 0.00 | 0.00 | 0.00 | 0.00 | 0.00 | 0.00 | 0.00 | 0.00 |
| 100.00 | 100.00 | 100.00 | 100.00 | 100.00 | 100.00 | 100.00 | 100.00 |
| 100.00 | 100.00 | 100.00 | 100.00 | 100.00 | 100.00 | 100.00 | 100.00 |
| 0.00 | 0.00 | 0.00 | 0.00 | 0.00 | 0.00 | 0.00 | 0.00 |
| 100.00 | 100.00 | 100.00 | 100.00 | 100.00 | 100.00 | 100.00 | 100.00 |
| 0.13 | 0.26 | 0.69 | 0.09 | 0.13 | 0.24 | 0.00 | 0.27 |
| 4.10 | 4.66 | 5.80 | 2.62 | 3.42 | 3.72 | 5.44 | 6.34 |
| 5.60 | 4.74 | 5.41 | 2.80 | 3.91 | 3.76 | 6.58 | 6.29 |
| 3.89 | 3.47 | 4.77 | 1.47 | 4.40 | 4.31 | 5.41 | 5.96 |

| PSI_SRR6966268 | PSI_SRR8429074 | PSI_SRR8429075 | PSI_SRR8429076 | PSI_SRR8429077 | PSI_SRR8429078 | PSI_SRR8429079 | PSI_SRR8429080 |
|---|---|---|---|---|---|---|---|
| /- | *Xrp1* +/- | | | *RpS12* G97D/G97D | | | *RpS1[illegible]* |
| 93.62 | 98.52 | 98.09 | 97.14 | 99.27 | 99.06 | 99.33 | 98.38 |
| 99.81 | 99.74 | 99.07 | 100.00 | 100.00 | 99.82 | 99.57 | 99.80 |
| 0.00 | 0.29 | 0.00 | 0.00 | 0.46 | 0.77 | 0.00 | 0.23 |
| 100.00 | 99.71 | 100.00 | 100.00 | 99.54 | 99.23 | 100.00 | 99.77 |
| 100.00 | 100.00 | 100.00 | 100.00 | 100.00 | 100.00 | 100.00 | 100.00 |
| 0.00 | 0.00 | 0.00 | 0.00 | 0.00 | 0.00 | 0.00 | 0.00 |
| 100.00 | 100.00 | 100.00 | 100.00 | 100.00 | 100.00 | 100.00 | 100.00 |
| 0.20 | 0.42 | 0.37 | 0.16 | 0.12 | 0.38 | 0.23 | 0.11 |
| 2.89 | 3.20 | 5.44 | 3.85 | 1.70 | 2.94 | 3.29 | 2.46 |
| 3.91 | 3.98 | 4.53 | 4.25 | 1.00 | 1.33 | 0.99 | 1.02 |
| 1.70 | 1.62 | 4.48 | 2.15 | 1.30 | 3.18 | 3.91 | 1.06 |

| PSI_SRR8429081 | PSI_SRR8429082 | MINPSI_ALL | MAXPSI_ALL |
|---|---|---|---|
| 2 *G97D/G97D* , *RpS3* *+/-* | | | |
| 99.52 | 99.09 | 90.87 | 99.52 |
| 100.00 | 99.52 | 99.07 | 100.00 |
| 0.23 | 0.00 | 0.00 | 0.77 |
| 99.77 | 100.00 | 99.23 | 100.00 |
| 100.00 | 100.00 | 100.00 | 100.00 |
| 0.00 | 0.00 | 0.00 | 0.00 |
| 100.00 | 100.00 | 100.00 | 100.00 |
| 0.00 | 0.00 | 0.00 | 0.69 |
| 3.05 | 4.40 | 1.70 | 6.34 |
| 2.12 | 2.55 | 0.99 | 6.58 |
| 2.25 | 3.27 | 1.06 | 5.96 |
