## Supplementary Table 2 for "RpS12-mediated induction of the Xrp1^short^ isoform links ribosomal protein mutations to cell competition"

**Supplemental Table 2:**

| **Genotypes** | **Heat-shock** | **N number** |
|---|---|---|
| **Figure 1A:** y w p{hs:FLP}/+; UAS-RpS12-Flag /+; act>stop>gal4, UAS- GFP/ Xrp1-HA | 25 min heat shock | **12** |
| **Figures 1B-1C:** y w p{hs:FLP}/+; UAS-RpS12-Flag /+; act>stop>gal4, UAS-GFP/+ | 25 min heat shock | **10** |
| **Figure 1D:** y w p{hs:FLP}/+; UAS-RpS12-Flag /UAS-Xrp1-RNAi ; act>stop>gal4, UAS- GFP/+ | 25 min heat shock | **8** |
| **Figure 1E:** nubGal4, UAS-RFP/UAS-RpS12-Flag; Xrp1-HA/+ | **no** | **12** |
| **Figure 1F:** nubGal4, UAS-RFP/ UAS-RpS12$^{G97D}$-Flag; Xrp1-HA/ + | **no** | **10** |
| **Figure 1G:** nubGal4, UAS-RFP/ UAS –RpL13-Flag; Xrp1-HA/ + | **no** | **5** |
| **Figure 1H:** nubGal4, UAS-RFP/ UAS –RpS17; Xrp1-HA/ + | **no** | **5** |
| **Figure 1I:** y w p{hs:FLP}/+; UAS-RpL13-Flag /+; act>stop>gal4, UAS-GFP/+ | 25 min heat shock | **5** |
| **Figure 1J:** y w p{hs:FLP}/+; UAS-RpS12$^{G97D}$-Flag /+ ; act>stop>gal4, UAS-GFP/+ | 25 min heat shock | **10** |
| **Figures 2B-D:**<br>RNAseq genotypes published previously (PMID: 30078730 and PMID:31841522)<br>WT genotype: w$^{11-18}$/+; FRT82B/+<br>RpS17$^{+/-}$ genotype: w$^{11-18}$/y w p{hs:FLP}; RpS17 p{ubi:GFP} FRT80B/+<br>RpS3$^{+/-}$ genotype:w$^{11-18}$/y w p{hs:FLP}; FRT82 RpS3 p{arm:LacZ}/+<br>Xrp1$^{+/-}$, RpS3$^{+/-}$ genotype: w$^{11-18}$/y w p{hs:FLP}; FRT82 RpS3 p{arm:LacZ}/FRT82B Xrp1M2-73<br>Xrp1$^{+/-}$ genotype: w$^{11-18}$/y w p{hs:FLP}; FRT82B Xrp1M2-73 /+<br>RpS12$^{G97D/G97D}$ genotype: w$^{11-18}$; rpS12 $^{G97D}$ FRT80B / rpS12 $^{G97D}$ FRT80B,<br>RpS12$^{G97D/G97D}$ ; RpS3$^{+/-}$ genotype: w$^{11-18}$; rpS12$^{G97D}$ FRT80B / rpS12 $^{G97D}$ FRT80B RpS3 | **no** | 3 biological samples for each genotype |
| **Figure 3A:**<br>Genotypes of Lane 1,2,4,5,6,7 and 9 same as RNA-seq genotypes in Figure 2.<br>Lane 1: WT: w$^{11-18}$/+; FRT82B/+<br>Lane 2: w$^{11-18}$/y w p{hs:FLP}; FRT82B Xrp1M2-73 /+<br>Lane 3: y w p{hs:FLP};Xrp1$^{08}$/Xrp1$^{08}$,<br>Lane 4: RpS12$^{G97D/G97D}$ genotype: w$^{11-18}$; rpS12 $^{G97D}$ FRT80B / rpS12 $^{G97D}$ FRT80B<br>Lane 5: RpS17$^{+/-}$ genotype: w$^{11-18}$/y w p{hs:FLP}; RpS17 | **no** | **N=3** |

| | | |
|---|---|---|
| p{ubi:GFP} FRT80B/+<br>Lane 6: RpS3$^{+/-}$ genotype:w$^{11-18}$/y w p{hs:FLP}; FRT82 RpS3 p{arm:LacZ}/+<br>Lane 7: Xrp1$^{+/-}$, RpS3$^{+/-}$ genotype: w$^{11-18}$/y w p{hs:FLP}; FRT82 RpS3 p{arm:LacZ}/FRT82B Xrp1M2-73<br>Lane 8: y w p{hs:FLP}; Xrp1$^{08}$ RpS3 /Xrp1$^{08}$,<br>Lane 9: RpS12$^{G97D/G97D}$ ; RpS3$^{+/-}$ genotype: w$^{11-18}$; rpS12$^{G97D}$ FRT80B / rpS12 $^{G97D}$ FRT80B RpS3<br>Lane 10: act>Gal4, UAS-CD8GFP/+ ,<br>Lane 11: act>Gal4, UAS-CD8GFP/UAS-RpS12-Flag ,<br>Lane 12: act>Gal4, UAS-CD8GFP/UAS-RpS12$^{G97D}$-Flag | | |
| **Figure 3C-D:**<br>Genotypes same as in Figure 2B-2D | **no** | |
| **Figure 3E:**<br>-act>Gal4, UAS-CD8-GFP/+ ,<br>-act>Gal4, UAS-CD8GFP/UAS-RpS12-Flag<br>- act>Gal4, UAS-CD8GFP/UAS-RpS12$^{G97D}$-Flag | **no** | **N=3** |
| **Figures 3F-G:**<br>- y w p{hs:FLP}; FRT82B/+,<br>- y w p{hs:FLP}; Xrp1$^{08}$/Xrp1$^{08}$,<br>- y w p{hs:FLP}; RpS3/+<br>- y w p{hs:FLP}; Xrp1$^{08}$ RpS3 /Xrp1$^{08}$, | **no** | **N=3** |
| **Figure 4A:**<br>nubGal4, UAS-RFP/UAS-Xrp1$^{short}$; Xrp1-HA/+, | **no** | **N=4** |
| **Figure 4B:**<br>y w p{hs:FLP}/+; UAS-Xrp1$^{short}$ /+; act>stop>gal4, UAS-GFP/ + | 25 min heat shock | **N=8** |
| **Figures 4C, D, E:**<br>y w p{hs:FLP}/+; UAS-Xrp1$^{short}$/act>stop>gal4, UAS-CD8-GFP+ | 45 min heat shock | **N=6** |
| **Figure 4F:**<br>nubGal4, UAS-RFP/UAS-Xrp1$^{short}$; Xrp1-HA/+, | **no** | **N=3** |
| **Figures 4G-H:**<br>y w p{hs:FLP}/+; act>stop>gal4, UAS-CD8-GFP/ UAS-Xrp1$^{short}$; Xrp1$^{KO/KO}$ | **4G:** 45 min heat shock<br>**4H:** 25 min heat shock | **N=8** |
| **Figure 4I:**<br>y w p{hs:FLP}/+; act>stop>gal4, UAS-CD8-GFP/ UAS-Xrp1$^{Long}$; Xrp1$^{KO/KO}$ | 45 min heat shock | **N=4** |
| **Figure 5A:**<br>y w p{hs:FLP}/ y w p{hs:FLP}; arm-lacZ M67c FRT80 Xrp1$^{Exlong}$/FRT80 Xrp1$^{Exlong}$ | 60 min heat shock | **N=3** |
| **Figure 5B:**<br>y w p{hs:FLP}/ y w p{hs:FLP}; arm-lacZ M67c FRT80 /FRT80 | 60 min heat shock | **N=6** |

| | | |
|---|---|---|
| **Figures 6A-D:**<br>Wild type genotype: $w^{11-18}$/+; FRT82B/+<br>$RpS3^{+/-}$ genotype:$w^{11-18}$/y w p{hs:FLP}; FRT82 RpS3 p{arm:LacZ}/+<br>$Xrp1^{+/-}$, $RpS3^{+/-}$ genotype: $w^{11-18}$/y w p{hs:FLP}; FRT82 RpS3 p{arm:LacZ}/FRT82B Xrp1M2-73 | **no** | **Biological replicates /(technical replicates)**<br>**Wild-type: 4/ (3)**<br>**RpS3+/-: 3/(3)**<br>**Xrp1+/-, RpS3+/-: 4/(3)** |
| **Figure 7A:** enGal4, UAS-GFP/ +; Xrp1-HA/UAS-SypRNAi (v33012) | **no** | **N=6** |
| **Figure 7B:** enGal4, UAS-GFP, $RpS18^{M56(f)}$/ +; Xrp1-HA/UAS-SypRNAi (v33012) | **no** | **N=6** |
| **Figure 7C:** enGal4, UAS-GFP/ UAS-SypRNAi (v110542); Xrp1-HA/+ | **no** | **N=5** |
| **Figure 7D:** enGal4, UAS-GFP, $RpS18^{M56(f)}$/ UAS-$Syp^{RNAi}$ (v110542); Xrp1-HA/+ | **no** | **N=6** |
| **Figure 7E:** y w p{hs:FLP}; UAS-$Syp^{RNAi}$ (v110542)/+; act>STOP>Gal4, UAS-GFP | 45 min heat shock | **N=6** |
| **Figure 7F:** y w p{hs:FLP}; UAS-$Syp^{RNAi}$ (v110542)/UAS-$Xrp1^{RNAi}$; act>STOP>Gal4, UAS-GFP | 45 min heat shock | **N=6** |
| **Figure S1A:**<br>nubGal4, UAS-RFP/UAS-RpL14; Xrp1-HA/+, | **no** | **N=10** |
| **Figure S1B:**<br>y w p{hs:FLP}/+; UAS-RpS17/+; act>stop>gal4, UAS- GFP/ + | 25 min heat shock | **N=5** |
| **Figure S1C:**<br>y w p{hs:FLP}/+; UAS-RpL14/+; act>stop>gal4, UAS- GFP/ + | 25 min heat shock | **N=5** |
| **Figure S1D:**<br>y w p{hs:FLP}/+; UAS-GFP-RpL10Ab/+; act>stop>gal4, UAS- GFP/ + | 25 min heat shock | **N=7** |
| **Figure S1E:**<br>enGal4, UAS-GFP/UAS-RpS12-Flag; Xrp1-HA/+ | **no** | **N=10** |
| **Figure S1F:**<br>enGal4, UAS-GFP/UAS-GFP-RpL10Ab; Xrp1-HA/+ | **no** | **N=5** |
| **Figures S2A-C and E:**<br>Genotypes same as in Figure 2B-2D | **no** | **See 2B-2D** |
| **Figures S3B and S3C:**<br>Genotypes same as in Figure 2B-2D | **no** | **See 2B-2D** |
| **Figure S3D:**<br>$RpS3^{+/-}$ genotype:$w^{11-18}$/y w p{hs:FLP}; FRT82 RpS3 p{arm:LacZ}/+<br>wild-type genotype: $w^{11-18}$/ y w p{hs:FLP}; FRT82B/+<br>$Xrp1^{+/-}$, $RpS3^{+/-}$ genotype: $w^{11-18}$/y w p{hs:FLP}; FRT82 RpS3 p{arm:LacZ}/FRT82B Xrp1M2-73 | **no** | **Wild-type: 2**<br>**RpS3+/-:2**<br>**Xrp1+/-, RpS3:1** |

| | | |
|---|---|---|
| **Figure S4A:**<br>nubGal4, UAS-RFP/UAS-Xrp1$^{long}$; Xrp1-HA/+, | **no** | **N=4** |
| **Figure S4B:**<br>nubGal4, UAS-RFP/+; Xrp1-HA/+, | **no** | **N=10** |
| **Figure S4C:**<br>y w p{hs:FLP}/+; UAS-Xrp1$^{long}$ /+; act>stop>gal4, UAS-GFP/ + | 25 min heat shock | **N=5** |
| **Figures S4D-S4G:**<br>y w p{hs:FLP}/+; UAS-Xrp1$^{short}$/act>stop>gal4, UAS-CD8-GFP | 45 min heat shock | **N=Same as 4C** |
| **Figure S4H:**<br>y w p{hs:FLP}/+; act>stop>gal4, UAS-CD8-GFP/+ | 45 min heat shock | **5** |
| **Figure S4I:**<br>nubGal4, UAS-RFP/UAS-Xrp1$^{long}$; Xrp1-HA/+ | **no** | **4** |
| **Figure S4J:**<br>nubGal4, UAS-RFP/+; Xrp1-HA/+, | **no** | **10** |
| **Figure S4K:**<br>y w p{hs:FLP}/+; act>stop>gal4, UAS-CD8-GFP/ UAS-Xrp1$^{short}$; Xrp1$^{KO/KO}$ | 45 min heat shock | **N=Same as 4G** |
| **Figure S4L:**<br>y w p{hs:FLP}/+; act>stop>gal4, UAS-CD8-GFP/ UAS-Xrp1$^{Long}$; Xrp1$^{KO/KO}$ | 45 min heat shock | **N=6** |
| **Figure S5A:**<br>y w p{hs:FLP}/ y w p{hs:FLP}; arm-lacZ M67c FRT80 Xrp1$^{Exlong}$/FRT80 Xrp1$^{Exlong}$ | 60 min heat shock | **N=3** |
| **Figure S6A:**<br>enGal4, UAS-GFP/ +; Xrp1-HA/UAS-wRNAi (BL 33623) | **no** | **N=8** |
| **Figure S6B:**<br>enGal4, UAS-GFP, *RpS18*$^{M56(f)}$/ +; Xrp1-HA/UAS- wRNAi (BL 33623) | **no** | **N=10** |
| **Figure S6C:**<br>y w p{hs:FLP}; act>STOP>Gal4, UAS-GFP/ UAS-Syp$^{RNAi}$ (v33012) | 25 min heat shock | **N=5** |
